# The whole-genome panorama of cancer drivers

**DOI:** 10.1101/190330

**Authors:** Radhakrishnan Sabarinathan, Oriol Pich, Iñigo Martincorena, Carlota Rubio-Perez, Malene Juul, Jeremiah Wala, Steven Schumacher, Ofer Shapira, Nikos Sidiropoulos, Sebastian M. Waszak, David Tamborero, Loris Mularoni, Esther Rheinbay, Henrik Hornshøj, Jordi Deu-Pons, Ferran Muiños, Johanna Bertl, Qianyun Guo, Chad J. Creighton, Joachim Weischenfeldt, Jan O. Korbel, Gad Getz, Peter J. Campbell, Jakob S. Pedersen, Rameen Beroukhim, Abel Gonzalez-Perez, Núria López-Bigas, on behalf of the PCAWG Drivers and Functional Interpretation Group and the ICGC/TCGA Pan-Cancer Analysis of Whole Genomes Network

## Abstract

The advance of personalized cancer medicine requires the accurate identification of the mutations driving each patient’s tumor. However, to date, we have only been able to obtain partial insights into the contribution of genomic events to tumor development. Here, we design a comprehensive approach to identify the driver mutations in each patient’s tumor and obtain a whole-genome panorama of driver events across more than 2,500 tumors from 37 types of cancer. This panorama includes coding and non-coding point mutations, copy number alterations and other genomic rearrangements of somatic origin, and potentially predisposing germline variants. We demonstrate that genomic events are at the root of virtually all tumors, with each carrying on average 4.6 driver events. Most individual tumors harbor a unique combination of drivers, and we uncover the most frequent co-occurring driver events. Half of all cancer genes are affected by several types of driver mutations. In summary, the panorama described here provides answers to fundamental questions in cancer genomics and bridges the gap between cancer genomics and personalized cancer medicine.

---

Tumors arise from genomic mutations, often referred to as ‘drivers’, that confer proliferative advantage to a cell^1^. Various classes of mutations, such as somatic point mutations (single nucleotide variants, or SNVs, multinucleotide variants, or MNVs, and short insertions and deletions, or indels), somatic copy number alterations (SCNAs) and somatic balanced genomic rearrangements (SGRs)^2^, can drive tumorigenesis. In addition to somatic driver mutations, some cancer mutations are inherited in the germline. Identifying drivers from the myriad of ‘passenger’ mutations present in tumor genomes has been a major focus of cancer genomics over the last 20 years. Recently, methods aimed at identifying signals of positive selection in the pattern of tumor mutations, with the assumption that cancer development follows a Darwinian evolutionary process, have succeeded in detecting genes affected by driver mutations^3–9^.

Despite enormous advances in cancer genomics in recent years, only partial views of the contribution of genomic mutations to tumorigenesis have been obtained. As a result, fundamental questions about the emergence of tumors remain unanswered. It is unclear, for instance, whether tumors of all cancer types are caused primarily by genomic driver events, and the relative contributions of point mutations and structural variants (SVs), such as SCNAs and SGRs to the emergence of cancer remain unknown. Moreover, beyond point mutations in the promoters of *TERT*^10,11^ and a few other genes^12^ and some instances of promoter and enhancer hijacking^13–15^, we have only limited insight into the contribution of non-coding mutations to tumorigenesis. A key open question concerns the number and specific combinations of driver mutations required to make a somatic cell become malignant and how this differs across tumors.

The characterization of more than 2,500 tumors by the International Cancer Genomics Consortium (ICGC) and The Cancer Genome Atlas (TCGA) under the Pan-Cancer Analysis of Whole Genomes (PCAWG)^16^ initiative provides an unprecedented opportunity to obtain a comprehensive whole-genome view of driver events in cancer. Here, we set the goal of identifying all driver events (somatic point mutations, SCNAs, SGRs, and potentially predisposing germline variants) across the whole genome of each tumor in the PCAWG cohort. We call this list of driver events the per-patient panorama of driver mutations (or simply panorama). To obtain it, we designed an incremental approach (Fig. 1a; Methods and Suppl. Notes 1 and 2) that exploits the power of this cohort to discover novel drivers, both coding and non-coding, and the knowledge accumulated through decades of cancer genetics research^2^. We found driver mutations in virtually all tumors, thereby providing definitive evidence of the oft-quoted maxim that cancer is fundamentally a genomics disease. We also demonstrated the presence of a small number of driver mutations in each tumor – 4.6 on average, a number that is relatively stable regardless of of the variability in mutational burden. While the contribution of somatic point mutations and SVs to tumorigenesis across cancer types differs, their relative proportions across the entire cohort are very similar. We found that most individual tumors harbor a unique combination of driver mutations, and we uncovered the most frequently co-occurring driver events. Some of these combinations may have a synergistic effect in the emergence of cancer, as in the cases of somatic point mutations of *KRAS* and *SMAD4* in pancreatic adenocarcinomas (Panc-AdenoCA) and *DAXX* and *MEN1* across pancreatic endocrine (Panc-Endocrine) tumors. Our analysis also revealed that up to 25% of tumors contain non-coding driver point mutations, the most frequent of which are *TERT* promoter mutations, with large differences between tumor types.

**Figure 1.**
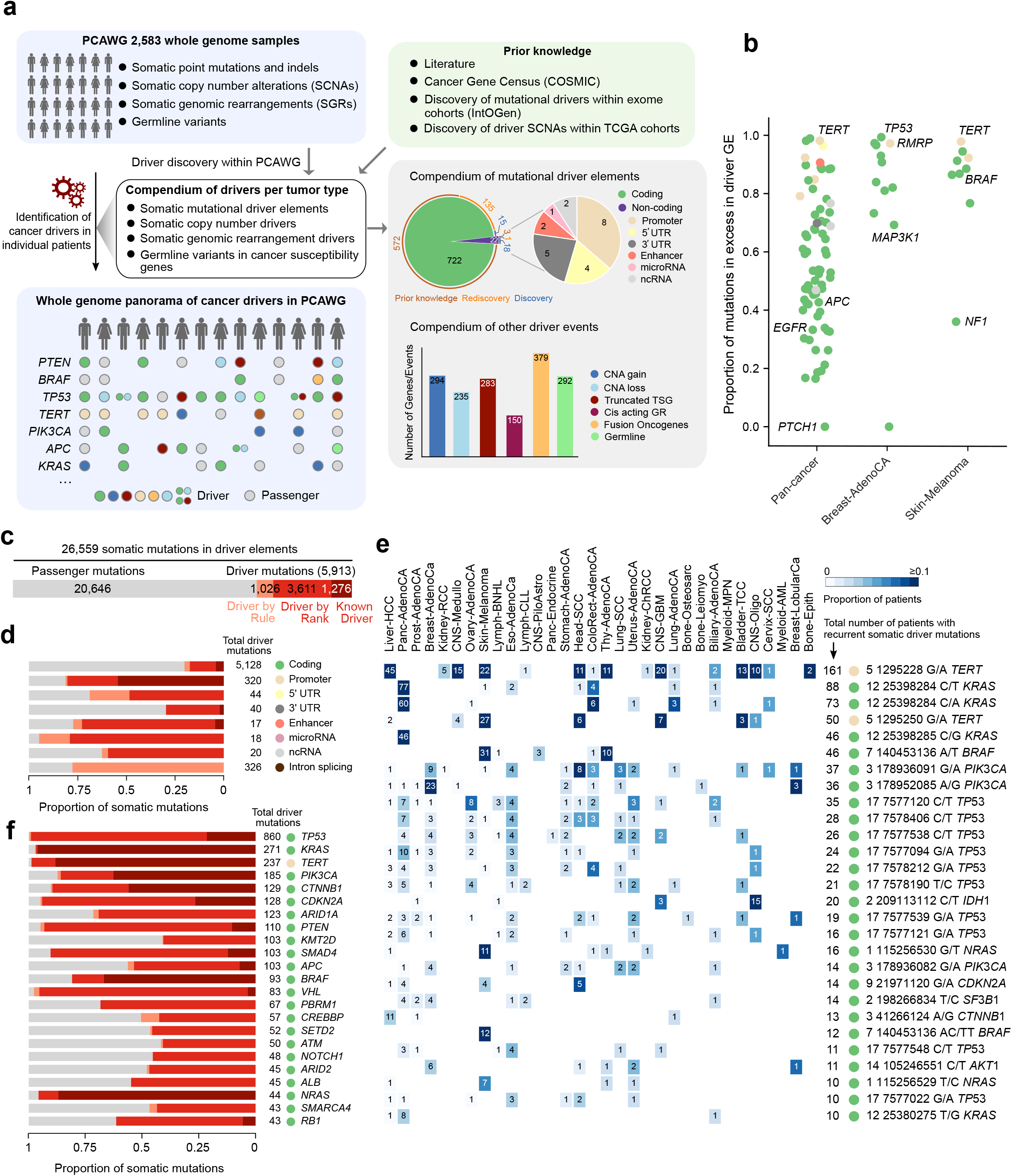
Approach to building a panorama of driver mutations across PCAWG tumors. (a) A schematic representation of the comprehensive incremental approach applied to identify all driver mutations in PCAWG tumors. GEs of different types with signals of positive selection in the cohort, and previously known driver GEs and point mutations obtained from other sources integrated the Mutational Compendium, which therefore comprises both novel discoveries and rediscoveries of drivers across the cohorts in PCAWG, color coded in the panel. Previously known and novel driver SVs and potentially tumorigenic germline variants were collected to build a Compendium of other driver events. (The list of tumor suppressor genes included in the search for SV drivers and susceptibility germline variants was obtained from the Mutational Compendium.) Using carefully calibrated methods we then employed these Compendia to identify the driver events across all tumors in the PCAWG cohort, thus producing the Whole genome panorama of cancer drivers in PCAWG. (b) Point mutations in discovery GEs may be either drivers or passengers; the fraction of the former is variable across GEs. It is computed as the excess of mutations above the background rate of each GE. The figure shows the distribution of the excess of mutations computed across GEs in the pan-cancer, breast adenocarcinoma and malignant melanoma cohorts. (Colors of GEs as in the Compendium in panel a). (c) The catalog of somatic driver point mutations comprises 5,913 driver point mutations, including Known Driver point mutations, driver point mutations identified via the rank-and-cut approach (Driver by Rank) and point mutations in other driver GEs identified via rules (Driver by Rule). The number of passenger mutations (in gray) in driver GEs is also shown. (d) The number and fraction of drivers among all point mutations identified varied across GEs of different nature. In the case of intronic point mutations affecting splice sites, only those affecting the canonical donor and acceptor sites and predicted to be loss-of-function in tumor suppressor genes by the LOFTEE^39^ are taken into account to compute the fraction of passengers. (e) Somatic driver point mutations in the cohort include both coding and non-coding events. The heatmap represents the recurrence of somatic driver point mutations across cohorts of GEs with at least 10 driver point mutations in PCAWG, with the color in each cell showing the proportion, and the number, the absolute count of mutated patients. The nature of the GEs affected by them is denoted by circles following the same color code as in Figure 1a. Note that each point mutation is defined by its position and nucleotide change; *KRAS* hotspot mutations caused by two different nucleotide changes appear at the second and third row. The columns of the heatmap are sorted according to the size of the cohort (high to low). (f) Identified driver point mutations in GEs that possess at least 40 across the cohort.

## Patient-centric catalog of somatic driver point mutations

We first focused on identifying somatic driver point mutations–SNVs, MNVs, and short indels– in each individual tumor. One necessary, but no sufficient step in this direction consists in the identification of cancer driver genomic elements in the cohort, a purpose for which in recent years several successful methods have been developed^4–9^. Using an array of these methods, the PCAWG Drivers and Functional Interpretation Group^17^ identified coding and non-coding genomic elements (GEs), including protein-coding genes, promoters, 5’UTRs, 3’UTRs, and enhancers with signals of positive selection across the PCAWG cohort (Fig. 1a; Methods). (Here, we refer to these drivers as discovery GEs.) While this reduces the problem, two important challenges remain.

First, not all somatic point mutations in these discovery GEs are driver mutations. This is revealed from the comparison of the computed expected rate of passenger point mutations (background) and the observed excess of point mutations above it-henceforth referred to as excess^8,18^ (Fig. 1b; Extended Data Fig. 1a). This excess constitutes the fraction of somatic driver point mutations of each GE in the cohort. For example, while virtually all somatic point mutations of *TP53* across breast adenocarcinomas (Breast-AdenoCA) are likely to be drivers (~100% computed excess), the excess computed for *NF1* somatic point mutations in malignant melanomas (Skin-Melanoma) indicates that about two thirds are likely passengers. Thus, to solve the aforementioned first problem, we designed a rank-and-cut approach to identify the somatic driver point mutations in each driver GE. Briefly, somatic point mutations observed across the cohort were ranked on the basis of a number of features that distinguish known tumorigenic mutations from passenger events. Then, top-ranking instances–up to the number computed from the excess– were nominated as drivers (Details in Suppl. Note 1 and Extended Data Figs. 1b-e).

Second, the statistical power of the PCAWG cohort to discover driver GEs is limited; some *bona fide* cancer genes could be mutated at frequencies below the detection threshold of current driver discovery methods^17,19^. To overcome this problem, we first collected GEs missed by the driver discovery process in PCAWG, which have been detected in other cohorts of tumor whole-exomes^20,21^, or identified through clinical or experimental evidence in recent decades^2^ (Fig. 1a). We call the integrated list of discovery and prior knowledge GEs, the Compendium of Mutational Driver GEs (Mutational Compendium, for short); Suppl. Table 1. Then, we designed a set of stringent rules to nominate somatic driver point mutations affecting the prior knowledge GEs on the basis of their known or inferred mode of action (Suppl. Note 1, Extended Data Fig. 1b and f). Somatic driver point mutations in those elements are assumed to be few, as may be deduced from their lack of signals of positive selection in the cohort. The decision-making process integrated by the rank-and-cut and the rule-based approaches, which we call onCohortDrive, accurately distinguishes between known driver and benign somatic point mutations and closely agrees with the estimated average number of driver coding mutations across cohorts using a dNdS approach^8^ (Suppl. Note 1; Extended Data Figs. 1g and 2a).

After applying onCohortDrive to all tumors in PCAWG, we obtained a patient-centric catalog comprising 5,913 somatic driver point mutations (Fig. 1c,d; Extended Data Fig. 2b; Suppl. Table 2). Interestingly, the single most recurrent somatic driver point mutation in the cohort affects the promoter of *TERT*, followed by coding SNVs affecting *KRAS, BRAF*, and *PIK3CA* (Fig. 1e). The coding sequence of *TP53* is the GE affected by the largest number of somatic driver point mutations in the cohort (n=860), with the promoter region of *TERT* ranking third (Fig. 1f). Interestingly, *TERT* is the most recurrently mutated driver GE in some cohorts, such as malignant melanomas (Skin-Melanoma) and glioblastomas (CNS-GBM) (Extended Data Fig. 2c).

## The role of somatic point mutations in tumorigenesis

We then used this catalog to answer open questions about the role of somatic point mutations in tumorigenesis. First, leveraging the unique opportunity provided by the comprehensive whole-genome view of drivers provided by PCAWG, we asked what is the relative contribution of somatic non-coding mutations to cancer.

The 785 somatic driver point mutations identified in non-coding regions –30% of which are *TERT* promoter point mutations and 41% intronic point mutations affecting the splice-sites of loss-of-function driver genes (Supp. Note 1)– constitute 13% of the catalog (Fig. 1d and Fig. 2b; table below the barplot). While this contribution may be underestimated because our detection of somatic driver point mutations is biased towards coding genes due to the use of prior knowledge, we obtain a similar result (12%) if the calculation is done only on discovery GEs (Suppl. Note 1; Extended Data Fig. 2d). Up to 25% of all PCAWG tumors contain at least one non-coding somatic driver point mutation (Fig. 2a). Interestingly, more than 50% of patients of certain types of tumors, such as bladder transitional cell carcinomas (Bladder-TCC) and glioblastomas (CNS-GBM), have at least one non-coding somatic driver point mutation. At the other extreme of the spectrum, in tumor types such as acute myeloid leukemias (Myeloid-AML) and pilocytic astrocytomas (CNS-PiloAstro), non-coding somatic driver point mutations appear to contribute little to tumorigenesis.

**Figure 2.**
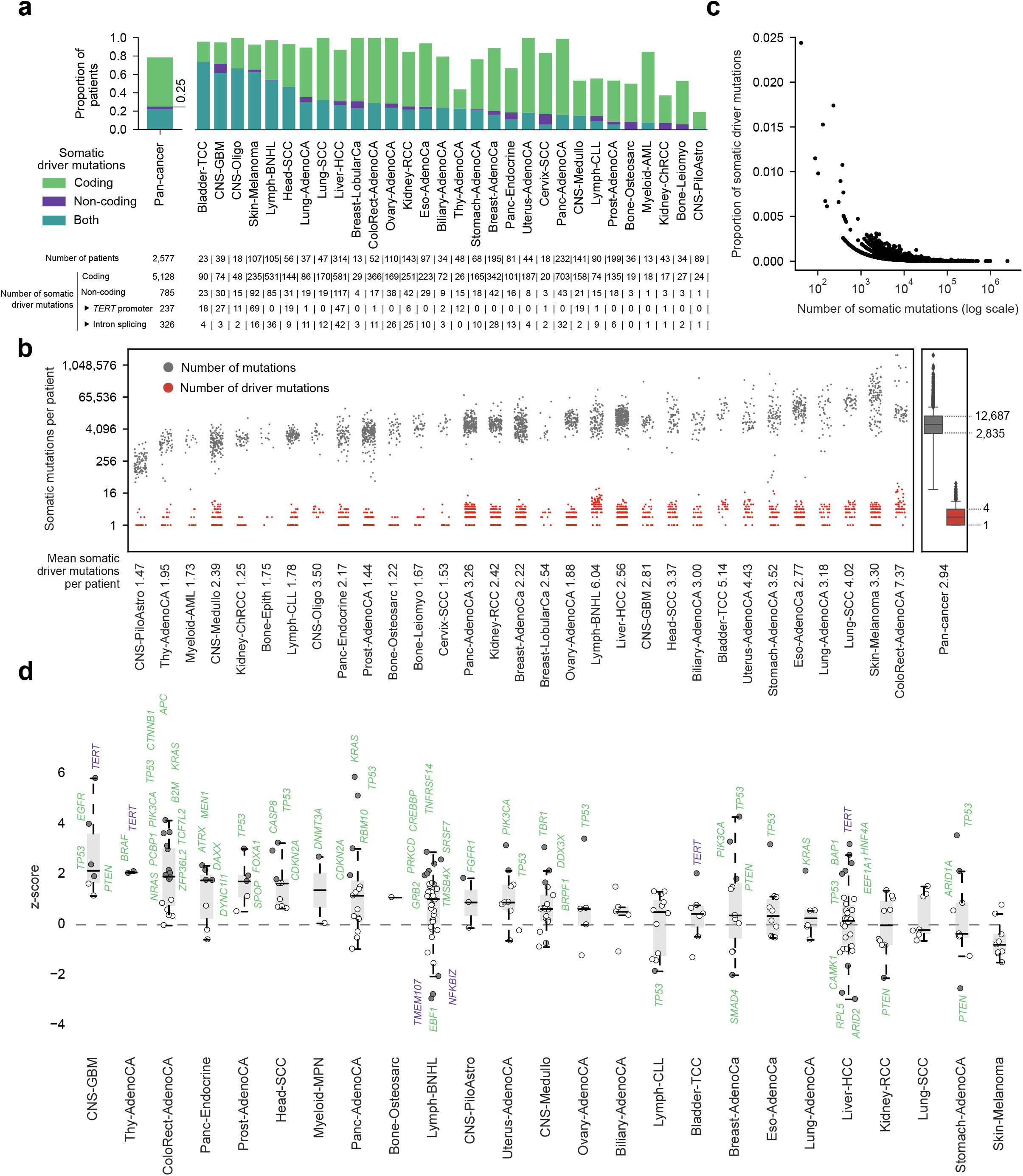
Catalog of somatic driver point mutations. (a) Number of tumors in each cohort and across all PCAWG bearing either coding (green), non-coding (blue), or both (purple) types of somatic driver point mutations. Below the graph, we present the number of patients, and the number of coding and non-coding driver point mutations (and specifically the number of *TERT* promoter and intron-splicing affecting point mutations in loss-of-function driver genes) in each cohort. (b) Tumors in the cohort exhibit mutational burden with a wide range of variation (gray dots and boxplot), whereas the number of driver point mutations remains remarkably stable (red dots and boxplot). (Only tumors with at least one somatic driver point mutation are included in the plot.) (c) The fraction of somatic driver point mutations in tumors and the total number of somatic point mutations are negatively correlated across the PCAWG cohort. (d) Normalized (z-score) distribution of the cancer cell fraction of GEs across tumor types. Filled circle represent GEs with point mutations cancer cell fraction deviating significantly from expectations (two-tail p-value < 0.05). The names of significant coding and non-coding GEs are colored following the convention in panel (a).

Next, we asked how many somatic driver point mutations occur in each tumor, and how this number varies with the mutational burden. We found that 78% of PCAWG tumors have at least one somatic driver point mutation, with a mean of 2.9 per tumor (Figs. 2b and 3b). This number is close to the average number of coding non-silent point mutations in excess recently estimated within genes in the Cancer Gene Census (CGC) across 7,664 TCGA tumors using a dNdS approach^8^. Strikingly, even heavily mutated tumors, such as malignant melanomas (Skin-Melanoma) and colorectal adenocarcinomas (ColoRect-AdenoCA), carry a mean of 3.3 and 7.4 driver point mutations, respectively. Overall, if the entire range of variation of the mutational burden across the cohort is considered (i.e., between just 41 and 2.5 million point mutations per genome), the number of driver point mutations remains remarkably stable, with the interquartile range between 1 and 4 (Fig. 2b). The number of driver point mutations increases more slowly than the total number of point mutations in PCAWG tumors (Extended Data Fig. 3a). As a result, the proportion of driver point mutations is anticorrelated with the mutational burden of tumors (Fig. 2c). Across most cancer types, driver point mutations show a higher cancer cell fraction than passenger point mutations (Extended Data Fig. 3b). This observation is more apparent for certain GEs in the Mutational Compendium, which exhibit a significant accumulation of highly clonal point mutations, thereby indicating that they carry potential founder or potent driver mutations^22^ (Fig. 2d).

In at least 6.7% (37 out of 554) of the tumors where we fail to identify somatic driver point mutations the cause may be an insufficient coverage in the sequencing (Extended Data Fig. 3c; PCAWG-Drivers Discovery). This affects in particular some cohorts, such as prostate adenocarcinomas (Prost-AdenoCA; 8 out of 93 patients with no somatic driver point mutations), stomach adenocarcinomas (Stomach-AdenoCA; 10 out of 16 patients), and Biliary adenocarcinomas (Biliary-AdenoCA; 5 out of 7 patients). The failure in other cohorts may be caused by the incompleteness of the Mutational Compendium or the prevalence of other types of genomic alterations.

In summary, despite large differences in mutational burden, tumors carry a stable low number of driver point mutations –2.9 as average across PCAWG. While non-coding point mutations represent only a small fraction (3%) amongst all point mutations in GEs of the Mutational Compendium, they contribute to tumorigenesis in 25% of the cohort, and constitute 13% of all driver point mutations in the PCAWG cohort.

## Integrated panorama of whole-genome driver mutations

In addition to somatic point mutations, tumors may be driven by somatic SVs, such as SCNAs, and SGRs. As with somatic point mutations, we first created a Compendium of somatic driver SVs (Fig. 1a), and then used strict rules to identify which SCNAs (Supp. Note 2; Extended Data Fig. 4a) and SGRs (Extended Data Fig. 4b, Methods) most likely act as drivers in each PCAWG tumor. Furthermore, we identified germline variants across the cohort which may also contribute to tumorigenesis, and revealed the landscape of biallelic inactivation events of tumor suppressor genes. The integration of the catalogs of somatic driver point mutations, SVs and germline variants across the cohort thus produced the first pan-cancer whole-genome panorama of driver events (Fig. 1a; Fig. 3a and b; Suppl. Table 3).

**Figure 3.**
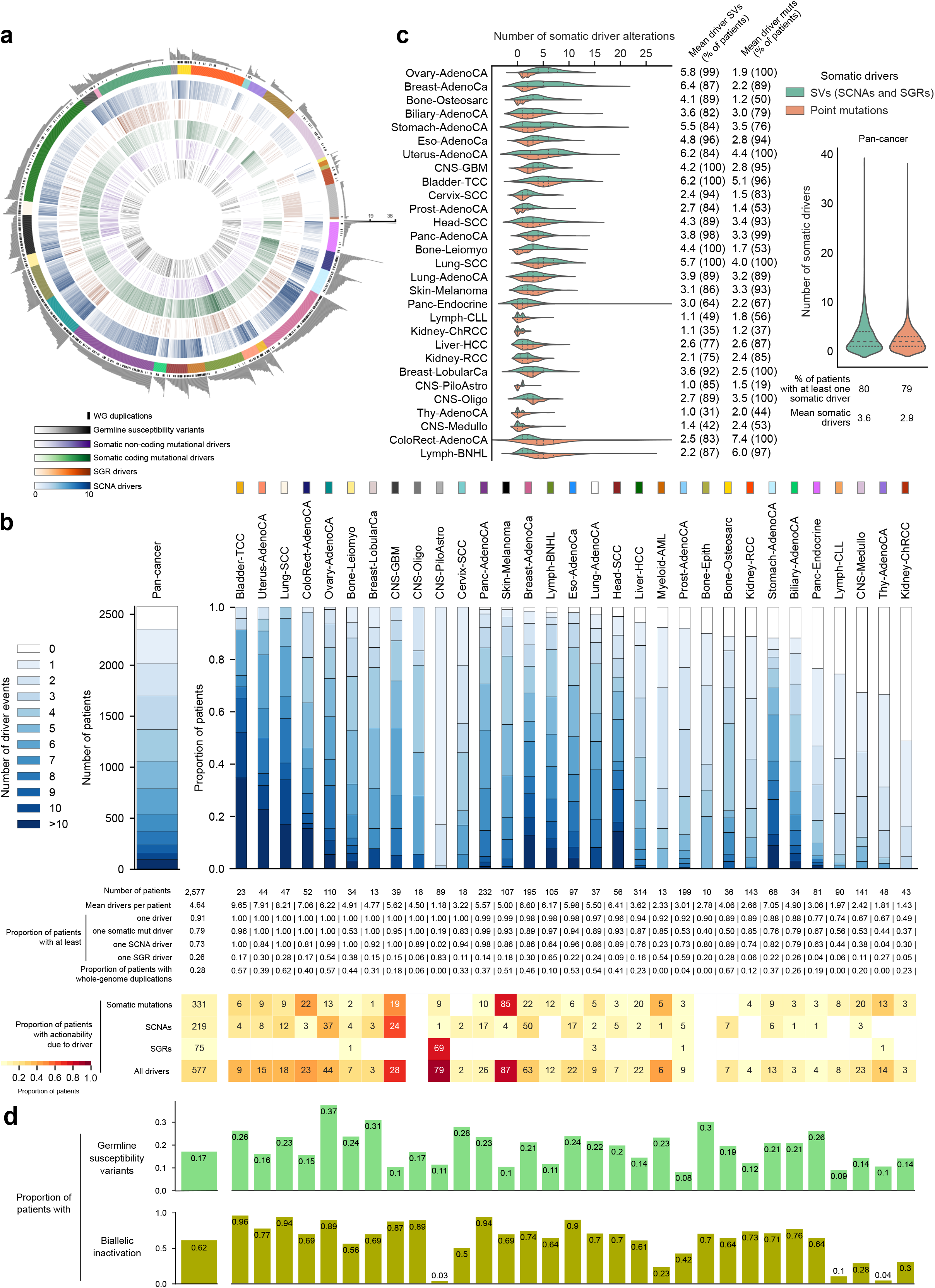
Whole-genome panorama of driver mutations. (a) The whole-genome panorama of driver mutations in the PCAWG cohort, represented as a circos plot. Each arc represents a tumor in the cohort. From the periphery to the center of the plot the layers represent: i) the total number of driver alterations in the tumor; ii) whether the patient bears a whole genome duplication; iii) the tumor type, with color codes specified in panel b; iv) the number of SCNAs; v) the number of SGRs; vi) somatic coding point mutations; vii) somatic non-coding point mutations; and viii) germline variants. (b) Row-stacked bar plot of the total number of tumors in each cohort with different counts of driver mutations. The table below the bar plot offers a complementary description of the panorama. The heatmap at the bottom represents the fraction (color) and number of tumors of each cancer type with driver point mutations, SCNAs, and SGRs that provide biomarkers of response to anti-cancer therapies either approved or under clinical trials. (c) Contribution of somatic point mutations and SVs to tumorigenesis across cancer types. (Split violin plots represent the smoothed probability distribution of the numbers of each type of mutations in each cohort, which expand beyond the boundaries of the actual distributions.) The halves of each violin plot represent the distribution of the total number of somatic driver point mutations (bottom, orange) and driver SVs (top, green) in the tumors of each cohort. (d) Fraction of tumors of each cohort affected by germline susceptibility variants (top), and biallelic inactivation events (bottom).

We asked whether the panorama gave support to the widespread notion that all tumors have a genomic origin. We found driver mutations in 91% of the tumors of the cohort, with variable fractions across tumor types (Fig. 3b). Of note, 17% of all PCAWG tumors carried at least one potentially tumorigenic germline variant (Fig. 3d). Our failure to detect genomic drivers in some cohorts, such as kidney chromophobe adenocarcinomas (Kidney-ChRCC) may be attributed to our incomplete knowledge of cancer genes in these malignancies or to a prevalence of epigenomic alterations. Nevertheless, our results demonstrate, to the extent allowed by current analyses, that genomic mutations drive at least 9 out of 10 tumors.

We then turned to the longstanding question of the minimum number of driver mutations needed to turn a cell malignant^23–26^. We found on average 4.6 driver mutations across tumors of the cohort, again with large differences across cancer types (table below Fig. 3b). Even tumors of intensively studied types of cancer – such as breast adenocarcinomas (Breast-AdenoCa)– for which accumulated prior knowledge probably encompasses most possible driver events, contain on average fewer than 7 driver mutations. If we assume that driver mutations accumulate over time during the development of cancer^22^, the number required to initiate a tumor is probably even lower. In agreement with this idea, we found that tumors diagnosed at older ages tend to carry more driver mutations (Extended Data Fig. 5a), although this general trend masks important differences among malignancies (Extended Data Fig. 5b and c).

Since there have been contradictory reports of the relative contribution of somatic SVs and somatic point mutations to tumorigenesis^27,28^, we next explored the relative prevalence of both types of alterations in the cohort. We discovered that both appear to play a role of similar preponderance, with a mean of 3.6 somatic driver SVs in 80% of patients with at least one somatic driver SV, and mean of 2.9 somatic driver point mutations in 79% of patients with at least one somatic driver point mutation (Fig. 3c). However, these numbers hide important differences across cancer types. For example, a higher contribution to tumorigenesis is made by somatic SVs in ovarian adenocarcinomas (Ovary-AdenoCA; mean of 5.8 SVs vs 1.9 point mutations) and breast adenocarcinomas (Breast-AdenoCa; mean of 6.4 SVs vs 2.2 point mutations). On the other hand, somatic point mutations make a larger contribution to colorectal adenocarcinomas (ColoRect-AdenoCA; mean of 2.5 SVs vs 7.4 point mutations) and mature B-cell lymphomas (Lymph-BNHL; mean of 2.2 SVs vs 6 point mutations). A major proportion of the patients with somatic driver mutations (1,735/2,577) harbor both types of drivers, whereas a small (and comparable) proportion of patients harbor either somatic driver point mutations (n=288) or somatic driver SVs (n=318) (Extended Data Fig. 4c). Furthermore, as with somatic point mutations, the number of driver SCNAs in tumors is stable, irrespective of the fraction of the genome involved in focal amplifications or deletions (Extended Data Fig. 6).

We next asked how frequent these biallelic hits –considering both germline and somatic mutations– affecting tumor suppressor genes^23^, are across tumors. We found instances of biallelic inactivation of tumor suppressor genes in more than half (62%) of PCAWG tumors (Fig. 3d). In some cohorts, such as bladder transitional cell carcinomas (Bladder-TCC), lung squamous cell adenocarcinomas (Lung-SCC) and Pancreatic adenocarcinomas (Panc-AdenoCA), more than 90% of tumors carry at least one biallelic hit, comprising either somatic/somatic or somatic/germline events.

We also found that chromosomal arm-level gains and losses and whole genome duplications appear to be a pervasive type of mutation in the PCAWG cohort (table below Fig. 3b; Extended Data Fig. 8). More than one fourth (28%) of the tumors suffer whole genome duplications, ranging between 0% of thyroid adenocarcinomas (Thy-AdenoCA), acute myeloid leukemia (Myeloid-AML), and chronic lymphocytic leukemia (Lymph-CLL), and 67% of bone osteosarcomas (Bone-Osteosarc). Although it is difficult to assess the role of chromosomal arm-level and whole-genome duplication events in tumorigenesis, their pervasiveness suggests that they contribute to this process.

We also examined the contribution of each type of driver mutation to the landscape of potential therapeutic actionability of the cohort (Fig. 3b, bottom panel). We matched drugs employed in the clinic –both approved or under clinical trials– to the panorama of driver mutations in PCAWG, using the Cancer Biomarkers Database^29^. We considered only mutations that constitute biomarkers of response to an anti-cancer drug in tumors bearing no concomitant biomarker of resistance to the same drug (Methods). Overall, 331 tumors presented actionable driver point mutations, 219 presented driver SCNAs, whereas only 75 tumors have actionable driver SGRs. Overall, 577 tumors in the cohort bear mutations that constitute biomarkers of response to drugs that are either FDA-approved or in clinical trials. Overall, 1,678 tumors in the cohort present drivers that may be matched to anti-cancer therapies either approved or in clinical trials either through guidelines or repurposing. Moreover 307 other tumors bear driver mutations susceptible of targeting using pre-clincal molecules (Extended Data Fig. 7). Interestingly, several elements with non-coding driver mutations contribute repurposing opportunities to this actionability landscape. These include mutations causing over-expression of *TERT*, which are susceptible to targeting via Eribulin, currently in pre-clinical trials for gliomas^30^. Moreover, considerable fractions of the tumors of some cohorts (183 across all PCAWG), given their mutation burden, could potentially benefit from immune checkpoint inhibitors^31^, either alone of in combination with targeted therapies (see Methods).

## GEs affected by different types of driver mutations

Next, we used the whole-genome panorama of cancer driver mutations to explore different ways of alterations of cancer genes that result in tumorigenesis. Overall, 54% (197/366) of genes bearing somatic somatic point mutations in the coding sequence are susceptible to other types of driver mutations (Fig. 4a and b). *TP53*, for instance, carries the highest number of somatic driver point mutations in the cohort (954 patients), with more than 50% of the patients affected in certain tumor types, such as pancreatic (Panc-AdenoCA), breast (Breast-AdenoCa), and ovarian adenocarcinomas (Ovary-AdenoCA). It is also affected by SCNA losses, truncations due to SGRs, and intronic splicing somatic point mutations. The expression of *TP53* is significantly decreased (Mann-Whitney P<0.0001) in tumors with intronic splice-site somatic point mutations, SCNA loss, truncation or coding truncating somatic point mutations (via nonsense-mediated decay^32^), compared to those with the wild-type form (Fig. 4c). On the other hand, *CDKN2B, PIK3CA, CTNNB1, ERG, MCL1* and *CCND1* are mostly affected by only one type of mutation across tumor types (Fig. 4b).

**Figure 4.**
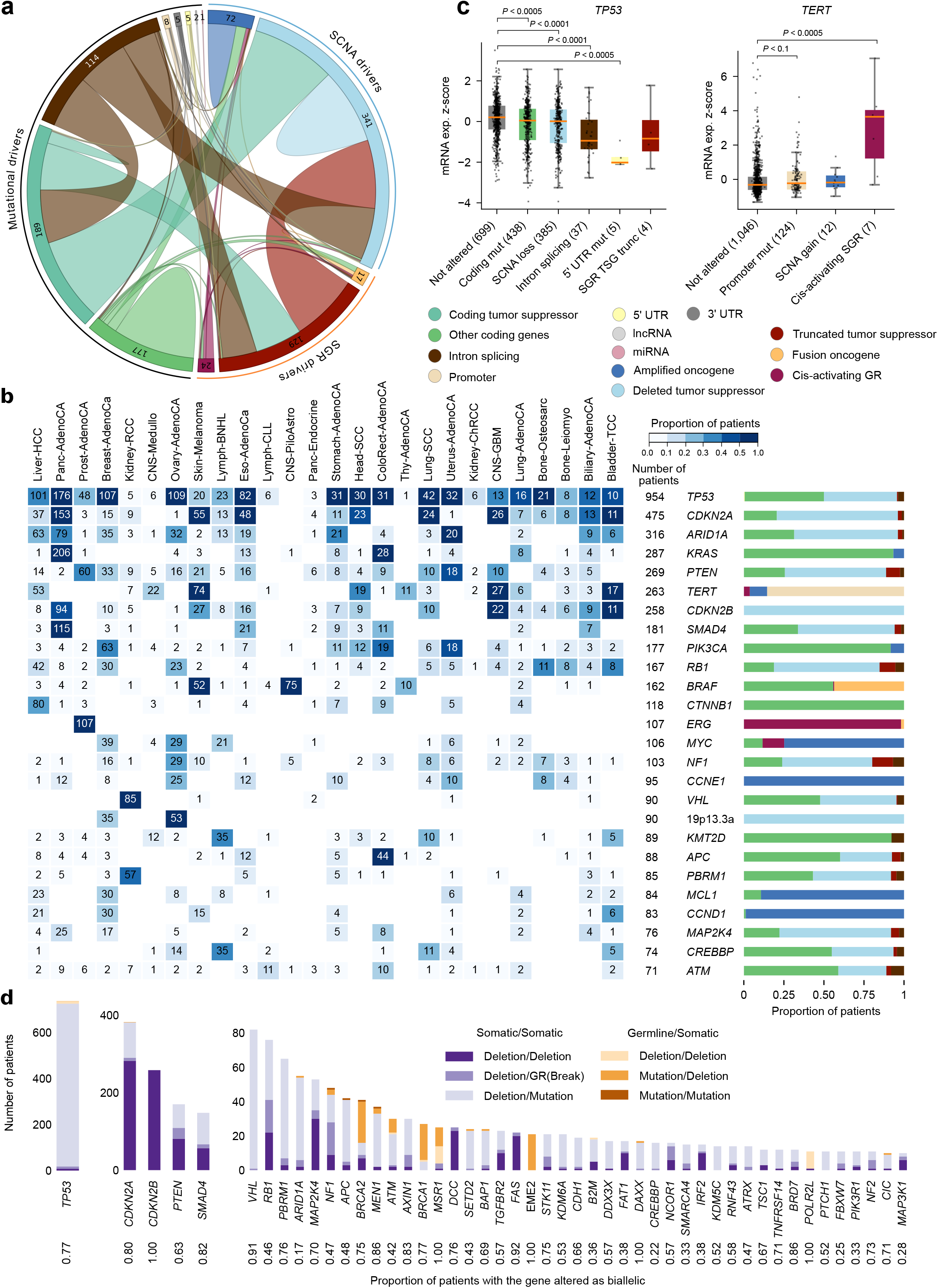
Multiple types of mutations affecting driver GEs. (a) Circos plot representation of the number of GEs in the Compendium affected by different types of driver mutations. Each arc in the circos represents GEs of a different nature; links between them depict GEs susceptible to two types of alterations, with their thickness proportional to the number of elements sharing both types of alterations. The numbers at the border of the plot represent unique counts of GEs suffering different types of mutations. (b) GEs targeted by different types of mutations in the cohort in more than 65 patients. The heatmap shows the recurrence of alterations experienced by each of them across cancer types (with the color indicating the proportion, and the number the absolute count of mutated patients); the barplot at the right reflects the proportion of each type of alteration affecting each GE. (c) Different types of driver mutations affecting a GE that result in the same type of change to their mRNA level. (d) Tumor suppressor genes showing biallelic inactivation in 10 or more patients.

*TERT* is the gene most frequently affected by non-coding somatic driver point mutations (Fig. 4b). It appears mutated in 263 patients through multiple routes, such as promoter SNVs and MNVs, balanced genomic rearrangements affecting cis-regulatory elements, and SCNA gains. *TERT* promoter somatic driver point mutations are particularly frequent in glioblastomas (CNS-GBM; 27/39), oligodendrogliomas (CNS-Oligo; 11/18), bladder transitional cell carcinomas (Bladder-TCC; 17/23), thyroid adenocarcinomas (Thy-AdenoCa; 11/48), and malignant melanomas (Skin-melanoma; 63/107). *TERT* SCNA gains are observed across malignant melanomas (Skin-melanoma; n=12), hepatocellular carcinomas (Liver-HCC; n=5), lung squamous (Lung-SCC; n=3) and adenocarcinomas (Lung-AdenoCa; n=5), whereas enhancer hijacking (due to SGRs) events affecting *TERT* are observed in malignant melanomas (Skin-melanoma; n=4), leiomyosarcomas (Bone-leiomyo; n=3), kidney renal clear cell (Kidney-RCC; n=1), kidney chromophobe renal clear cell (Kidney-ChRCC; n=1) and hepatocellular carcinomas (Liver-HCC; n=1). Notably, all these routes of driver mutation lead to the over-expression of *TERT* (Fig. 4c).

Finally, we studied the biallelic inactivation of tumor suppressor genes, that comprise combinations of germline and/or somatic driver events. Somatic/somatic events are more frequent (84%) among all genes with biallelic inactivation than germline/somatic events (32%) (Fig. 4d). The most frequent biallelically mutated gene, *TP53*, is inactivated across 736 tumors of 27 cancer types, most of which (707) harbor a somatic driver point mutation affecting one allele and the somatic loss of the other allele. Overall, we were able to detect the biallelic inactivation of *TP53* in 77% of the tumors with *TP53* driver mutations; in the remaining fraction, other mechanisms such as promoter hypermethylation may play a role. Well-known tumor suppressor genes, such as *CDKN2A* and *CDKN2B*, are rarely inactivated by point mutations, but rather appear homozygously lost (282 and 258 tumors, respectively). On the other hand, biallelic inactivation of *BRCA1* and *BRCA2* appears primarily due to one germline variant, followed by the somatic loss of the second allele in 63% (22/35 patients) and 45% (25/55 patients) of tumors with *BRCA1* and *BRCA2* driver mutations, respectively. The number of genes with biallelic inactivation in individual tumors differ within and across cancer types (Extended Data Fig. 9a). For example, in both bladder transitional cell carcinomas (Bladder-TCC) (18/23) and pancreatic adenocarcinomas (Panc-AdenoCA) (183/232), around 77% of patients carry more than one tumor suppressor gene with biallelic inactivation.

## Combinations of driver mutations

The panorama confirms the accepted notion that in most cases tumorigenesis is due to the accumulation of driver mutations rather than single driver events: 2,016 tumors (78%) of the cohort possess driver mutations affecting more than one GE. Moreover, almost each tumor is unique in their combination of driver events: among these 2,016 tumors there are 1,915 unique driver combinations. We then measured the overlap between the set of driver GEs of each pair of PCAWG tumors as their Jaccard index (Methods; Fig. 5a). As expected, tumors of the same cancer type tend to share a higher fraction of their drivers (diagonal of the lower triangular heatmap) than tumors of different cancer types. As a trend, CNS-Piloastro patients share the largest fraction of drivers relative to their total number (Jaccard distribution skewed to high values; mean Jaccard Index=0.62), followed by Panc-AdenoCA patients (0.3), Kidney-RCC (0.18) and ColoRect-AdenoCA (0.17). At the other end of the spectrum, Breast-AdenoCA (0.07), Liver-HCC (0.06), Stomach-AdenoCA (0.06) and CNS-Medullo (0.04) patients exhibit the largest degree of driver heterogeneity. The homogeneity of CNS-Piloastro, Panc-AdenoCA and Kidney-RCC may be explained at least in part by relatively few driver GEs identified in the cancer type and relatively few drivers per patient (diagonal of the upper triangular heatmap; Extended Data Fig. 10a). However, this is not the case for ColoRect-AdenoCA tumors, a case in which the relative homogeneity of drivers may stem from a rather narrow pathway to tumorigenesis, followed by genetic innovation after the establishment of the tumor. Conversely, the relative heterogeneity of drivers across CNS-Medullo, a cancer type with few driver GEs and few driver alterations per tumor may indicate the existence of multiple different pathways towards tumorigenesis.

**Figure 5.**
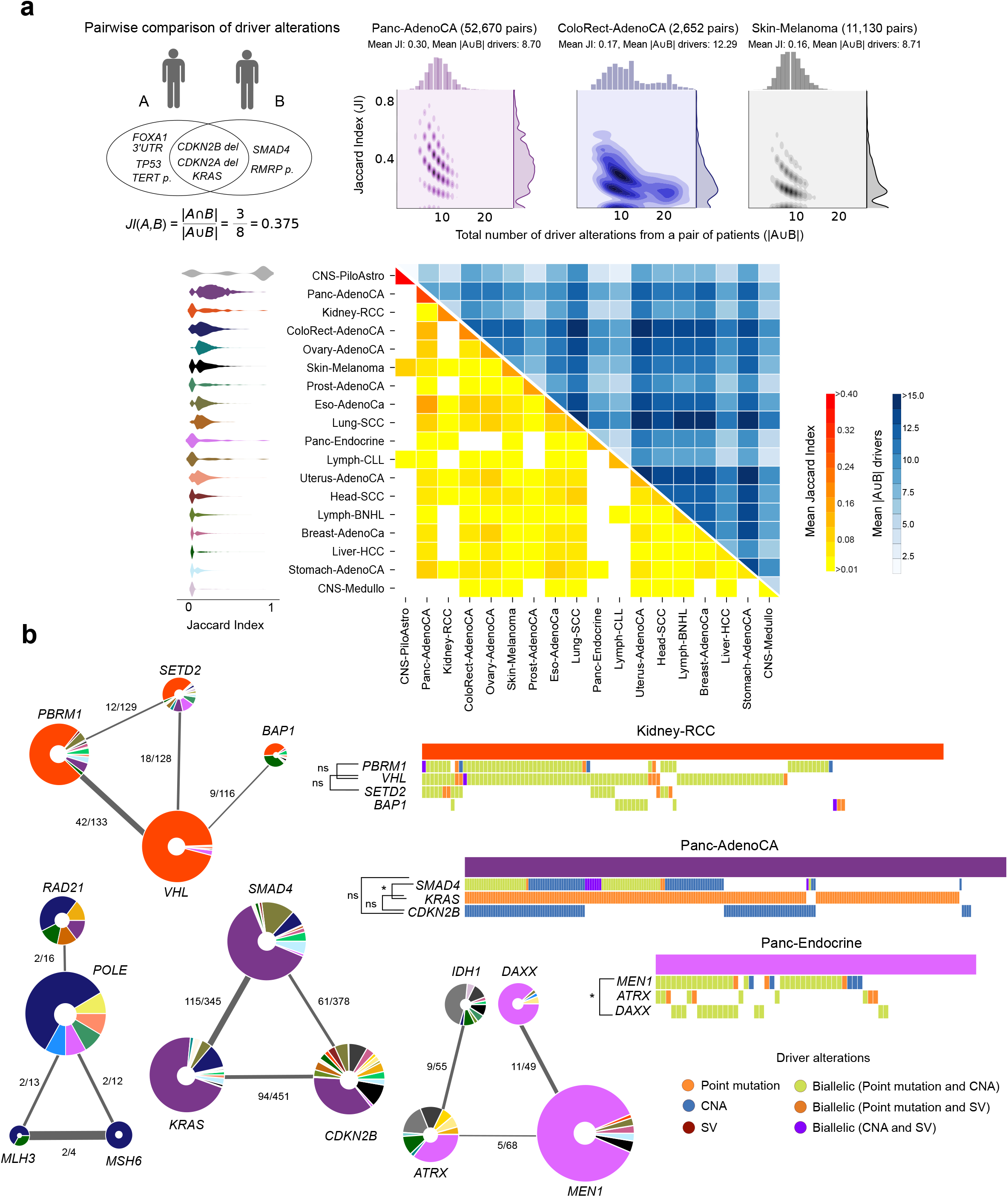
Combinations of driver mutations in the cohort. (a) For each pair of patients in the cohort we compute a Jaccard index of the overlap of their sets of driver mutations (at the level of driver GEs). The joinplots above the heatmap present the (smoothed) distributions of Jaccard index (right side) and number of patients in the union (top) and the 2D distribution of both variables across all pairs of patients of three selected tumor types. The mean Jaccard index of pairs of patients of each possible pair of tumor types are represented in the lower triangle heatmap. The violinplots represent the (smoothed) distribution of Jaccard index between pairs of patients of all cancer types in the heatmap. The mean number of drivers in the union of pair of patients of each possible pair of tumor types are represented in the upper triangle heatmap. Only pairs of patients of cancer types with at least 40 patients are included in the heatmap. (b) Selected groups of GEs whose driver mutations tend to co-occur across the cohort. Each donut plot in the graphs represent a GE, with the radius proportional to the number of patients carrying driver mutations affecting it, and the colors in the donut representing their recurrence in each cohort. The width of the links in the graphs are proportional to the Jaccard index of the overlap of patients with driver mutations affecting both GEs. The heatmaps at the right side represent the cohorts in which the highest recurrence of driver mutations (represented by the color tiles beside each gene name) affecting the members of each graph. Asterisk beside pairs of gene names indicate the significance of the observed co-ocurrence of somatic driver point mutations of several pairs of GEs are included in each heatmap.

We then searched for specific groups of drivers that tend to co-occur across tumors. To that end, we computed the Jaccard index of the patients in which each pair of driver GEs occur (Extended Data Fig. 10b). Figure 5b highlights several cases of highly co-occurring drivers. These include the already reported^33–35^ cooccurring biallelic inactivation of *VHL, PBRM1, SETD2* and *BAP1* across Kidney-RCC patients, the cooccurring biallelic inactivation of *MEN1* and *ATRX/DAXX* in Panc-Endocrine tumors, and the co-occurring biallelic or monoallelic inactivation of *SMAD4* and *CDKN2B* with the activating point mutations of *KRAS* in Panc-AdenoCA. The co-occurrence of point mutations involved in the biallelic inactivations of *VHL, PBRM1* and *SETD2* in Kidney-RCC tumors is not significant (P_VHL-pBRM1_=0.14, P_SETD2-VHL_=0.37), indicating this co-occurrence could be determined solely by the frequency of the biallelic inactivation of each driver GE. On the other hand, point mutations involved in the biallelic inactivation of *DAXX* and *MEN1* across Panc-Endocrine tumors co-occur more often than expected from their frequencies alone (P=0.03), as do activating point mutations of *KRAS* and mutations involved in the biallelic inactivation of *SMAD4* in Panc-AdenoCA patients (P=0.02). This suggests that there is at least some degree of synergy between them.

## Discussion

In the present study, we designed and implemented a systematic approach to comprehensively identify driver mutations in each tumor of the PCAWG cohort, with the final aim of exploring the genomic roots of tumorigenesis. Although thorough, this panorama of driver mutations is not complete because the approaches used to identify driver events possess limitations. Technical problems of the sequencing and the calling of variants may result in missing driver point mutations. The discovery of driver GEs is limited to regions of the genome with known functional elements, basically coding and non-coding genes and their cis-regulatory elements. Mutations affecting other regions of the genome and perturbing, for example, the higher-order structure of the chromatin may also contribute to tumorigenesis but will be missed by the methods designed to detect signals of positive selection in well-defined GEs. Also, some mutations affecting driver GEs which may still contribute to tumorigenesis at low frequency, may be overlooked by the drivers discovery until larger cohorts of whole-genomes become available^36^. The catalog of driver point mutations is incomplete with respect to non-coding driver mutations, due to the scarcity of prior knowledge of non-coding driver GEs. Furthermore, the accuracy in the estimation of the excess of mutations in GEs –in particular in noncoding regions– above their expected background may be affected by yet unknown mutational processes. This probably also affects the identification of driver SVs in some cohorts of yet unexplored cancer types, since only few events –due mostly to the lack of statistical power– were discovered in the PCAWG cohort^37^.

The panorama of driver mutations of PCAWG tumors reveals that almost all tumors contain genomic driver alterations (91% had at least one type of driver mutation), implying that the presence of somatic mutations is a condition necessary –although perhaps not sufficient^38^– for virtually all tumors. One fifth of the PCAWG tumors had at least one driver-balanced rearrangement, 73% at least one driver SCNA, 79% at least one driver point mutation, and 17% carry at least one germline susceptibility variant (Fig. 6). A prototypical glioblastoma (CNS-GBM), for example, bears a somatic driver point mutation activating the *TERT* promoter, the biallelic loss of *CDKN2A* and *CDKN2B*, the hyperactivation of EGFR via either a SCNA gain or a coding somatic point mutation, and/or the double somatic biallelic inactivation of *TP53*. The panorama also showed that the relative contribution of non-coding mutations is about 13%; this contribution surpasses 50% of the patients in some types of cancer. We also described that while overall somatic point mutations and SVs represent similar proportions of all driver events, this observation conceals large differences across cancer types, with some preponderantly harboring somatic driver point mutations and others driven primarily by somatic SVs. Another striking finding stemming from the panorama is that the variation in the number of somatic driver mutations required to initiate a tumor is not related to the total burden of mutations of the tumor. Instead, it is fairly constant and tends to be fewer than a dozen events in virtually all tumors. Finally, through the analysis of the sets of co-occurring driver mutations, the panorama provide a key piece in the efforts to understand how tumors originate and evolve.

**Figure 6.**
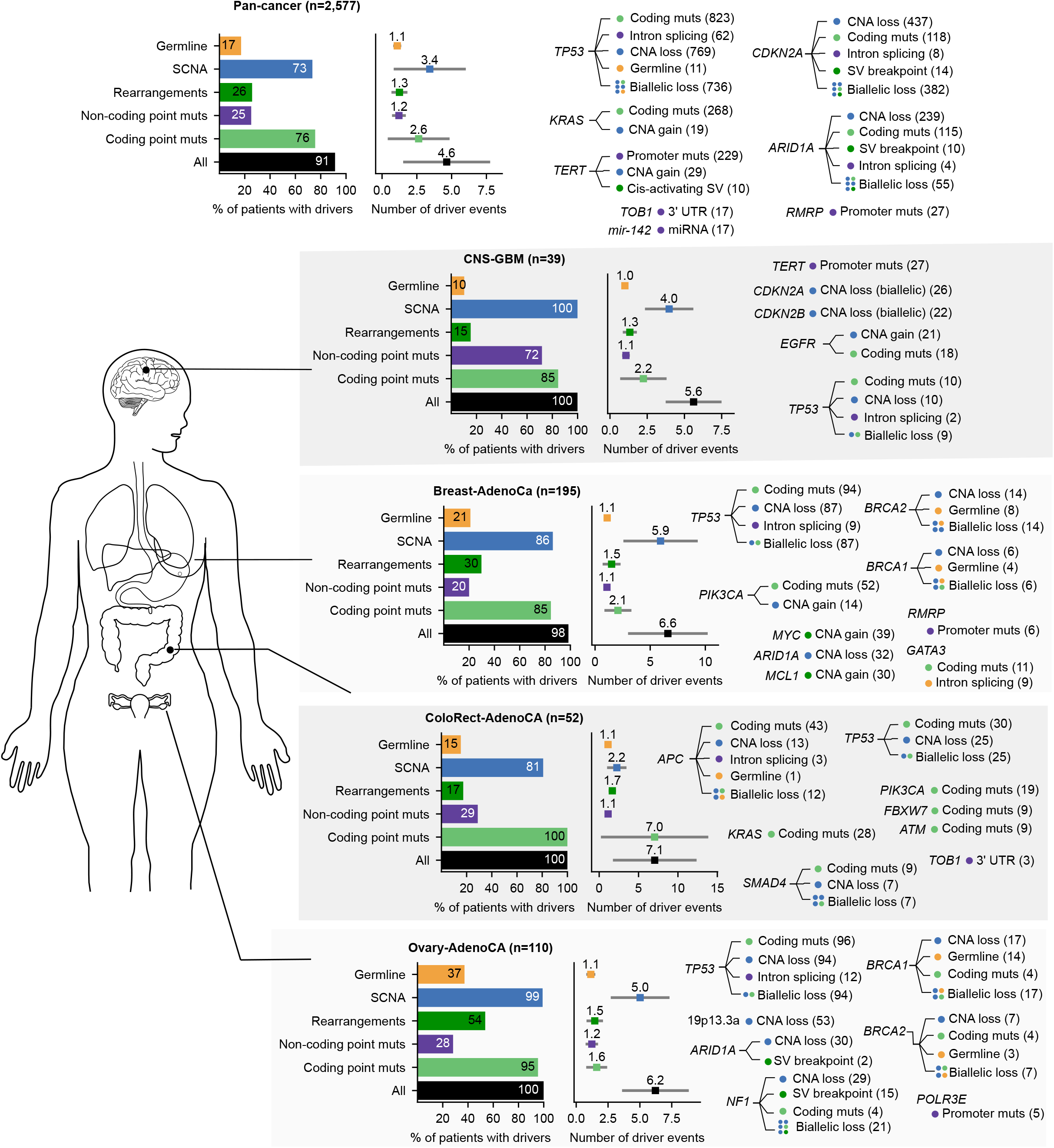
Snapshots of the panorama. Each row summarizes either the pan-cancer cohort or the cohort of a particular tumor type. The horizontal bar plot (at the left of each row) represents the proportion of patients with different types of mutations. The dot plot with whiskers at the right represents the mean number of each type of driver mutations registered across tumors with at least one event (the square dot), and its standard deviation (gray whiskers). For each tumor type, a representative set of genes which receive frequent driver mutations is presented at the right side of each row. For the full list of driver mutations see Supplementary Table 3.

The recent advances in cancer genomics described in the introduction have reliably identified genes driving tumorigenesis across cohorts of patients of different cancer types. However, the overarching objective of selecting the most effective treatments to target tumors –the personalized cancer medicine paradigm– requires the identification of the specific mutations driving the tumorigenesis in each patient, which remains a largely unsolved problem. The approach that we have described and implemented here to solve this challenge using the PCAWG cohort constitutes a proof-of-principle that bridges the gap between cancer genomics and personalized cancer medicine. We envision that personalized medicine initiatives, via systematic sequencing the tumor of cancer patients will make use of this type of approaches to support clinical decision-making.

## METHODS

### Genomics data

From the PCAWG Consortium^16^ we obtained the somatic point mutations (single and multiple nucleotide variants and short indels; syn7364923)^16^, copy number alterations (syn8042880)^37^, structural variants (syn7596712)^37^, cancer cell fraction (syn8532425)^22^, and expression data (syn3104297)^40^ across 2,583 tumors in the cohort. (Eight non-malignant samples were excluded from downstream analyses.) Genomic elements (GEs) that potentially harbor driver point mutations were obtained from the PCAWG Drivers and Functional Interpretation Group^17^, using the coordinates of Gencode v19 (syn5259890). The list of these genomic elements included the coding DNA sequence (CDS) of protein-coding genes, their 5’UTR and 3’UTR, the coordinates of their intronic acceptor and donor splice sites, their promoter regions, known enhancers, and the exonic regions of non-coding RNAs, and the promoter regions of the latter. In addition, we included *TERT* promoter mutations (chr5:1295228:G>A and chr5:1295250:G>A) in 80 samples, which were not detected in the PCAWG somatic point mutation calling due to low sequence coverage, in accordance with Sieverling *et al*. (2017)^41^.

### The Mutational Compendium and the Compendium of other driver GEs

We retrieved the list of mutational discovery GEs identified as drivers of PCAWG tumors from the PCAWG Drivers and Functional Interpretation Group^17^. This list of GEs covers coding genes, promoters, enhancers, 3’ and 5’ UTRs, and ncRNAs. Each GE in this list was detected because its mutational patterns exhibited signals of positive selection across one of several cohorts within PCAWG, the reunion of several cohorts of tumors with the same cell of origin or from the same organ (metacohorts), or across the entire cohort (pancancer cohort). They were therefore annotated as mutational drivers in each cohort, metacohort and/or the pan-cancer cohort. We also retrieved the excess of somatic mutations above their expected number for each GE in the respective cohorts, metacohorts and/or the pan-cancer cohort estimated by the NBR algorithm^8^. These are the discovery GEs. Five coding genes and one promoter region of a coding gene identified in the lymphoma cohorts with a high proportion of mutations introduced by AID (>40%) were removed from these discovery GEs.

In an effort to complement the list of discovery drivers and to add prior knowledge about the drivers of tumorigenesis to the compendium, we manually collated and curated a list of previously known cancer GEs – driving tumorigenesis through point mutations, SCNAs and/or balanced genomic rearrangements– across tumor types. This list of drivers comprises: a) GEs with validated tumorigenic effect, obtained from the CGC^2^ and literature reports (with special attention paid to recently reported instances of driver non-coding genomic elements), and b) genes whose mutational patterns show signals of positive selection across previously analyzed cohorts of tumor exomes or genomes^20,21^. To add SCNA driver GEs to the Discovery Compendium, we relied on the identification of amplified or deleted chromosomic regions (peaks) with significant recurrence across TCGA cohorts (see below). To add SGR driver events to the Compendium, we manually gathered known events from the literature and searched for their occurrence in the PCAWG cohort. Only rearrangements that appear in tumors of the cohort were included in the panorama (see below).

### Functional annotation of point mutations

The Variant Effect Predictor^39^ v86 was used with GRCh37 coordinates to determine the GEs affected by point mutations and their consequence types. The Combined Annotation Dependent Depletion (CADD) v. 1.0 score of all single and multiple nucleotide variants –as well as the maximum possible CADD score of short indels– was obtained from cadd.gs.washington.edu^42^.

### Identification of driver point mutations

We developed a method, onCohortDrive, to identify driver mutations affecting driver GEs in individual tumors. Supplementary Note 1 contains the full description of the method and its benchmarking. OnCohortDrive comprises a rank-based approach and a rule-based approach. The rank-based approach is applied to point mutations in GEs with an observed excess of point mutations above the number expected across the cohort. Briefly, consider that *N* is the total number of point mutations affecting a GE, and *n* is the number of point mutations in excess above the background rate across the tumors in a cohort, with *k* known driver point mutations. The *k* known driver point mutations are first nominated as drivers and withdrawn from the rank-based process. Then, the number of point mutations available to the rank-based approach is *N-k*, and the excess after identifying known tumorigenic point mutations is *n-k*. These *N-k* point mutations are then ranked, based on a combination of i) their functional impact scores, ii) whether they belong to mutational clusters, iii) their unlikeliness in the given tumor and iv) element specific features (e.g. the impact on TFBS for promoter and enhancers, the impact on secondary RNA structure for microRNAs, etc). OnCohortDrive then nominates as driver-by-rank the *n-k* top ranking mutations. Finally, the number of mutations in excess in the pan-cancer cohort after the excess in all cohorts has been processed (residual excess) are processed as explained above across all tumors in the pan-cancer cohort, excluding those in cohorts already processed.

Point mutations observed in prior knowledge GEs or in GEs with no mutational excess are not eligible for the rank-based approach and subsequently undergo a rule-based approach. The rules used to nominate driver-by-rule point mutations are based on the same features used for ranking. Point mutations that are not known drivers, drivers-by-rank, or drivers-by-rule are deemed passengers.

To make an unbiased assessment of the contribution of non-coding somatic point mutations to tumorigenesis, we applied onCohortDrive solely to somatic point mutations observed in discovery GEs across the PCAWG cohort, skipping the identification of known driver point mutations in GEs.

To evaluate the performance of onCohortDrive we carried out three benchmarks (described in full in Supplementary Note 1; Extended Data Fig. 1g). First, we assessed the accuracy of the approach (rank-and-cut and rule-based) to correctly classify as drivers and passengers 1,032 known cancer mutations and 12 known benign mutations in tumors in the PCAWG cohort from a total of 22,854 SNVs in coding genes of the Compendium. We found that while only 16% of the 22,854 SNVs are classified as drivers, 95% of known drivers (982 out of 1,032) are correctly classified. All 12 benign mutations are correctly classified as passengers. Second, to verify the stringency of the rules designed within onCohortDrive, we benchmarked the capacity of the rule-based approach to correctly classify the same set of driver and passenger point mutations. As expected, a lower proportion of SNVs are classified as drivers by rules, 2,412 (11%), still maintaining a high proportion of the known driver mutations correctly classified, 805 (78%), and all 12 benign variants classified correctly as passengers. Third, we evaluated the performance of the approach to correctly label as passengers all benign SNVs (ClinVar^43^; n=314) and a list of common coding polymorphisms obtained from (ExAC^44^; n=1,413) in genes of the Compendium. OncohortDrive correctly classifies 97% and 99% of those SNVs, respectively as passengers. Furthermore, we compared the number of coding driver point mutations identified by onCohortDrive across cohorts with the global average number of point mutations in excess estimated by the dNdScv^8^ algorithm in genes of the Compendium to carry out a sanity check of the catalog produced by the method. This revealed a satisfactory degree of agreement across cohorts (Supplementary Note 1; Extended Data Fig. 2a).

### Bias towards high CCF of point mutations in driver GEs

We computed the mean cancer cell fraction (CCF) of all point mutations affecting GEs identified across cancer types in PCAWG. We obtained the values of CCF of each somatic driver point mutation from the PCAWG Evolution and Heterogeneity Working Group^22^. To determine whether the mean CCF of a GE exhibited a significant bias towards high or low clonality, we randomly sampled the same number of mutations observed in each element from all the mutations in the same cancer type. By iterating this process 10,000 times, we computed an expected distribution of mean CCFs of each element and assessed whether a significant deviation from the observed mean existed.

### Identification of driver SCNAs

The Compendium of SCNA GEs was populated from the analysis of chromosomal regions that exhibit signals of positive selection in their patterns of focal amplifications or deletions across cohorts of tumors analyzed by TCGA. Significant SCNA peaks were detected by applying the GISTIC algorithm^45^ to both TCGA individual cohorts and to the pan-cancer cohort. Peaks detected in this manner were then merged into metapeaks and known driver genes within each metapeak were identified, when known. The list of metapeaks was further filtered by checking whether the expression of the driver contained within the metapeak exhibited a significant shift in coherence with the type of peak (amplification or deletion) in the same cohort in which it had been annotated as a driver. The metapeaks passing this filter integrated the Compendium of driver SCNAs. Finally, the copy number of the genomic segments overlapping with the loci of the metapeaks in the Compendium in PCAWG tumors was used to decide whether they were amplified, diploid or deleted. (See Supplementary Note 2 for details.)

### Identification of driver SGRs

The driver SGRs we considered included genic fusions involving an oncogene, truncation of tumor suppressors, and cis-activating rearrangements (e.g., promoter-rearrangement and enhancer hijacking). These events were obtained from literature reports, curated databases (Cancer Gene Census)^2^, and a set of high-confidence novel genomic rearrangements that were identified in the PCAWG cohorts (provided by PCAWG Structural Variants analysis working group). Using this information, each tumor within PCAWG was probed for the presence of driver rearrangements. In the case of gene fusions, the gene coordinates plus a 50-kb flanking window on either side of each member of the pair of fused genes were scanned for the presence of rearrangements. Furthermore, the rearrangements were annotated when they produced a sense in-frame fusion.

This resulted in 331 fusion events in 319 samples. For 214 of these events (in 204 samples) we could not find the expression evidence (i.e., expression of fusion transcripts) due to the lack of expression data (RNA-seq) available for those samples. However, for the 117 events (in 115 samples) with expression data available, we identified 41 events (in 40 samples) that have fusion transcript match, based on the results provided by PCAWG3 (syn10003873)^40^. For the remaining 76 fusion events (in 76 samples), the lack of fusion transcript match may be explained by promoter/enhancer hijacking events that resulted in an over-expression of the target oncogene. The majority of the fusion events that fall under this category are related to the fusion with IGH/IGL locus. On the other hand, we have included fusion events in seven samples based on the evidence from fusion transcripts (PCAWG3 results), but were not detected based on the aforementioned SV analysis. In addition, we have included four fusion events in nine samples of CNS-PiloAstro samples, based on a previous study^46^.

In the case of tumor suppressor genes, the breakpoints affecting exons were considered as drivers. In addition, we analyzed rearrangements affecting the cis-regulatory elements of the coding genes in the compendium. This included rearrangements in the promoter regions (promoter-SV) and those causing a juxtaposition of enhancers close to a gene (enhancer-hijacking). In the latter case, we focused on genes that CESAM analysis^13^ has shown to become over-expressed through the enhancer-hijacking process, which takes into account the breakpoints, SCNAs, target gene (mRNA) expression, and chromatin interaction data from topologically associating domains (TADs). For those genes, we performed CESAM analysis to identify PCAWG samples with genes that showed over-expression.

### Identification of likely tumorigenic germline variants

We identified all truncating (stop gain, frameshit, splice site) germline points mutations and rare germline SV deletions affecting genes within a cancer susceptibility list^47^. We also identified all truncating germline point mutations and SV deletions affecting genes involved in DNA repair^48^, given that a second inactivating event, either somatic (truncating or missense) or germline (truncating) was observed in the other allele.

### Identification of biallelic driver events

To identify tumor suppressor two-hit events^23^, we defined biallelic inactivation as a gene locus G^A/B^, where alleles A and B are genetically altered, leading to a genetic G^mut/mut^ state. The biallelic inactivation assessment includes three genetic inactivation event types consisting of somatic or germline deletions (“Loss”), somatic or germline SVs (“Break”) and somatic or germline SNVs (“Mutation”). Given a heterozygous G^A/B^ locus, we required a loss of the A allele of the gene, leading to a hemizygous G^−/B^ state, and genetic inactivation of the remaining B allele, specifically requiring the second event to overlap the loss on the A allele, leading to biallelic inactivation. We considered four classes of biallelic inactivations: i) Loss/Mutation, nonsynonymous driver mutations of the B allele; ii) Loss/Loss, two deletion events that overlap an exon and the copy-number derived allele count is 0 both for A and B allele; iii) Loss/Break, SVs where one or both breakpoints are situated in an exon of the B allele; and iv) Mutation/Mutation, a nonsynonymous germline SNV and a nonsynonymous driver somatic SNV of the same gene. We infer the germline mutation to occur on the A allele and the somatic mutation on the B allele, with the assumption that two independent driver mutation events are highly unlikely to occur on the same allele. All biallelic inactivation events involving at least one Loss event which had not been detected in the process of identification of driver SCNAs –either because the SCNA GE was under the statistical power of detection in the corresponding cohort, failed the expression filter, or because the Loss event involved an arm-level deletion (see Supplementary Note 2)– were included in the panorama as driver SCNA loss events.

### The panorama of driver mutations

All driver mutations identified across all tumors in the cohort (point mutations, SCNAs, and genomic rearrangements, as well as biallelic events, described in previous sections) were integrated at the tumor level to obtain the whole-genome panorama of driver events. This panorama is presented at the level of single patients in Figure 3a and summarized as tumor-type bar plot in Figure 3b. Throughout the paper we have used the panorama to answer open questions in cancer genomics, identify GEs affected by different types of driver events, and pairs or groups of driver mutations that either co-occur or are mutually exclusive across tumors. The panorama of driver mutations in the PCAWG tumors can be explored via prepared Gitools^49^ interactive heatmaps (http://www.gitools.org/pcawg) and browsed in IntOGen^20^, at http://www.intogen.org/pcawg (Supplementary Note 3).

### Gene expression changes caused by different types of driver alteration

To test the hypothesis that events affecting the same coding gene cause the same alteration to the expression of the gene –as in the case of promoter driver mutations, SCNAs, and SVs– we compared the expression of several genes (Zscore normalized across cohorts) in various sets of tumors with different genotypes via a Wilcoxon-Mann-Whitney test.

### Therapeutic landscape of PCAWG tumors

To match drugs employed in the clinic, both approved and under trials, as well as small molecules in pre-clinical research to the driver alterations in tumors of the cohort, we employed the Cancer Biomarkers Database and the Cancer Actionability Database^29^. We also considered the possibilities of repurposing drugs in any of these three groups to counteract a different alteration or to use in another tumor type. To compute the contribution of different types of mutations to the therapeutic landscape, we only accounted for matches that i) involve a drug approved for its use in the clinic or in clinical trials; ii) elicit sensitivity to the drug; iii) occur in the tumor type of the clinical guidelines specified for the biomarker.

### Combinations of drivers

For each pair of patients in the cohort, we computed the Jaccard index of the overlap of the driver GEs found affected in each as the ratio between the number of driver GEs in the intersection between the two samples and the number of unique driver GEs in both. Then for each pair of GEs in the Compendium, we did the same calculation with the overlap of the patients in which each was found to bear driver mutations. We assessed the statistical significance of the co-occurrence of somatic driver point mutations using a previously developed mutex algorithm^21^. The algorithm computes an empirical p-value through the comparison of the observed co-occurrence of mutations of the pair of genes with 10,000 randomly generated arrays of mutations takes into account the mutational frequency of each GE and the mutational burden of each tumor.

## Acknowledgments

NL-B acknowledges funding by the European Research Council (Consolidator Grant 682398). IRB Barcelona is a recipient of a Severo Ochoa Centre of Excellence Award from the Spanish Ministry of Economy and Competitiveness (MINECO; Government of Spain) and is supported by CERCA (Generalitat de Catalunya). R.S. is supported by an EMBO Long-Term Fellowship (ALTF 568-2014) co-funded by the European Commission (EMBOCOFUND2012, GA-2012-600394) support from Marie Curie Actions. A.G.-P. is supported by a Ramón y Cajal contract (RYC-2013-14554). O.P. is supported by the BIST PhD Fellowship. D.T. is supported by project SAF2015-74072-JIN, which is funded by the Agencia Estatal de Investigacion (AEI) and Fondo Europeo de Desarrollo Regional (FEDER). We thank Federico Abascal for his contribution to Extended Data Figure 2a. S.M.W. received funding through an SNSF Early Postdoc Mobility Fellowship (P2ELP3_155365) and an EMBO Long-Term Fellowship (ALTF 755-2014). The results published here are in part based upon data generated by the TCGA Research Network: http://cancergenome.nih.gov/. We thank, in particular, members of the Technical Working Group of PCAWG for their work in generating the somatic mutation calls used in this work.

## EXTENDED DATA FIGURE LEGENDS

**Extended Data Figure 1.**
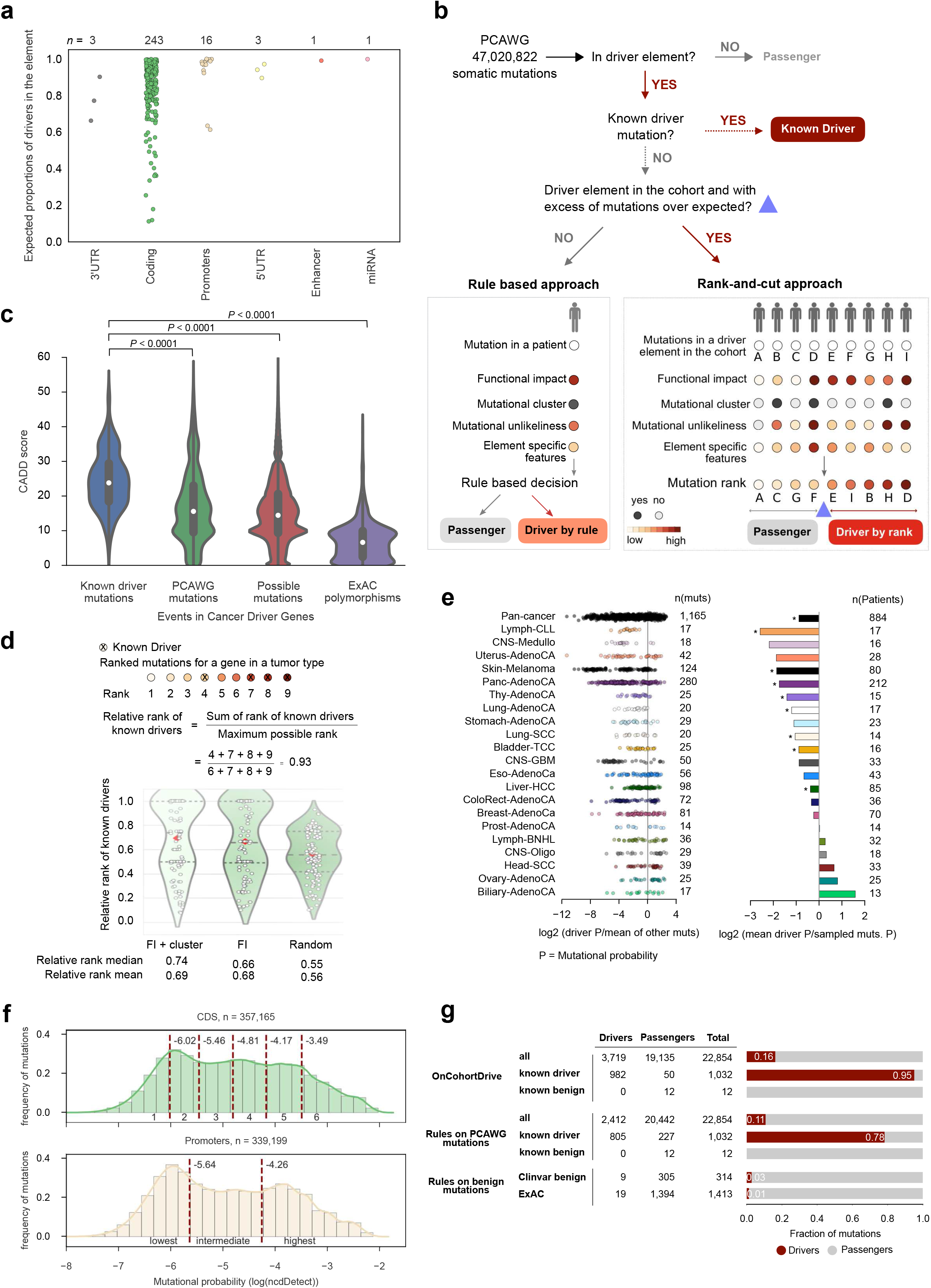
The onCohortDrive method. (a) Distribution of the expected proportion of drivers in each type of GE, computed using NBR (Supplementary Note 1) as the excess of mutations of each element above the background rate. (b) Point mutations in Genomic elements (GEs) follow different processes (either rank-and-cut approach or rule-based) to be nominated as drivers, depending on whether we are able to compute the excess of somatic mutations above the background rate in these GEs. In the rank-based process (right panel), mutations affecting each GE are ranked according to several features, and the rank is then cut at the position equal to the estimated excess of mutations in the element above the background rate. Mutations above the cut position are considered drivers, and those below, passengers. In the rule-based process (left panel), mutations in each tumor are nominated as drivers or passengers using the same features employed in the rank-based approach. (Details in Supplementary Note 1.) (c) Distribution of CADD scores of groups of variants in cancer genes. Sets of mutations, from left to right: i) known driver point mutations in cancer genes observed in PCAWG samples; ii) all other mutations in cancer genes in PCAWG; iii) all possible variants in cancer genes; iv) all polymorphisms observed in ExAC in cancer genes. The distributions of CADD scores of different sets of mutations were compared using the Kolmogorov-Smirnoff test. (Details in Supplementary Note 1.) (d) Relative rank of known driver mutations within all mutations observed in the PCAWG cohort. The violinplots represent the distribution, from right to left, of random relative ranking of known driver mutations, functional impact based ranking, and functional impact and clustering based ranking. (Details in Supplementary Note 1.) (e) Distribution of mutational probability (computed using ncdDetect) of known driver mutations relative to other mutations. Known driver mutations overall have significantly lower probability to occur than other mutations observed in tumors. In the left panel each dot is a known driver mutation in a patient and its probability of occurrence, relative to the probability of all other mutations in the patient is shown. The bars in the right panel show the comparison of the average probability of known driver mutations to the average probability of other sampled mutations (size equal to the number of known driver mutations) in each cohort. The cohorts with significantly (P<0.01) lower probability of occurrence of known mutations are highlighted (*). (Details in Supplementary Note 1.) (f) Distribution of the probability of occurrence of point mutations (ncdDetect) in coding genes (top, green) and promoters (bottom, yellow). The vertical broken lines mark the cutoffs of probability used to make the groups of point mutations considered separately in the rule-based approach. (g) Results of the benchmark of onCohortDrive on groups of known driver and benign mutations. The three main rows in the table describe the three datasets used in the benchmark (described in Supplementary Note 1), and the columns represent (from left to right) the number of variants in each dataset classified by onCohortDrive as drivers or passengers, and the total number of variants included in each group.

**Extended Data Figure 2.**
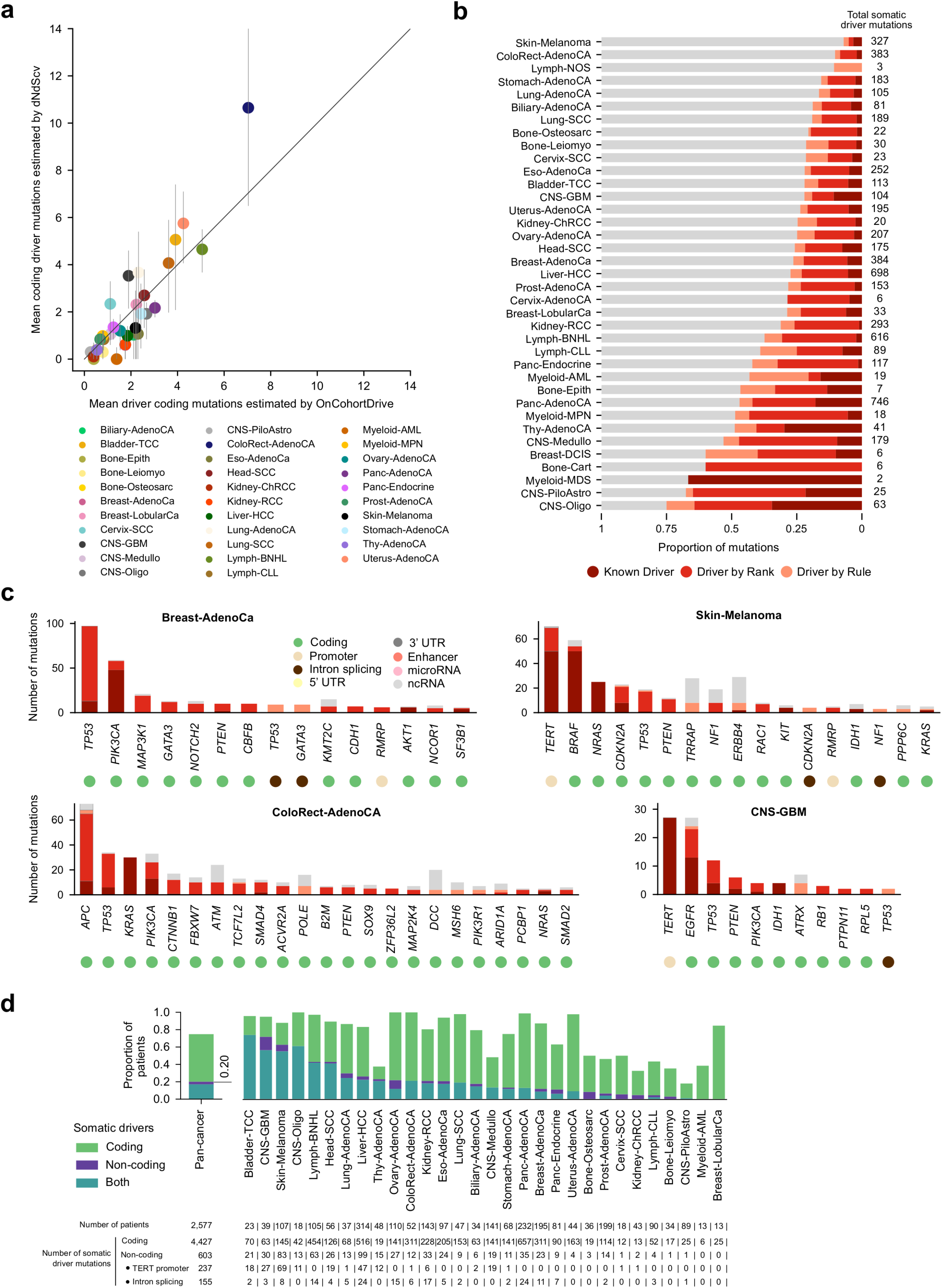
Building the catalog of driver point mutations. (a) Comparison between the average number of somatic coding driver point mutations identified by onCohortDrive across each cohort and the average (and confidence intervals) of mutations in excess identified in the same set of genes in the same tumors by the dNdSCV method. (b) Fraction of point mutations affecting GEs in different cancer types and in the pan-cancer cohort that are identified as known-driver, driver-by-rank, driver-by-rule in the catalog of driver point mutations. (Cohorts are sorted by the proportion of driver point mutations identified across each of them, and the classes of driver point mutations are color-coded as in Fig. 1c.) (c) Fraction of known-driver, driver-by-rank, driver-by-rule and passenger mutations in selected GEs across four cohorts in PCAWG. (d) Unbiased contribution of non-coding point mutations to tumorigenesis. Similar to main Figure 2a, but computed only from the discovery GEs (Supplementary Note 1)

**Extended Data Figure 3.**
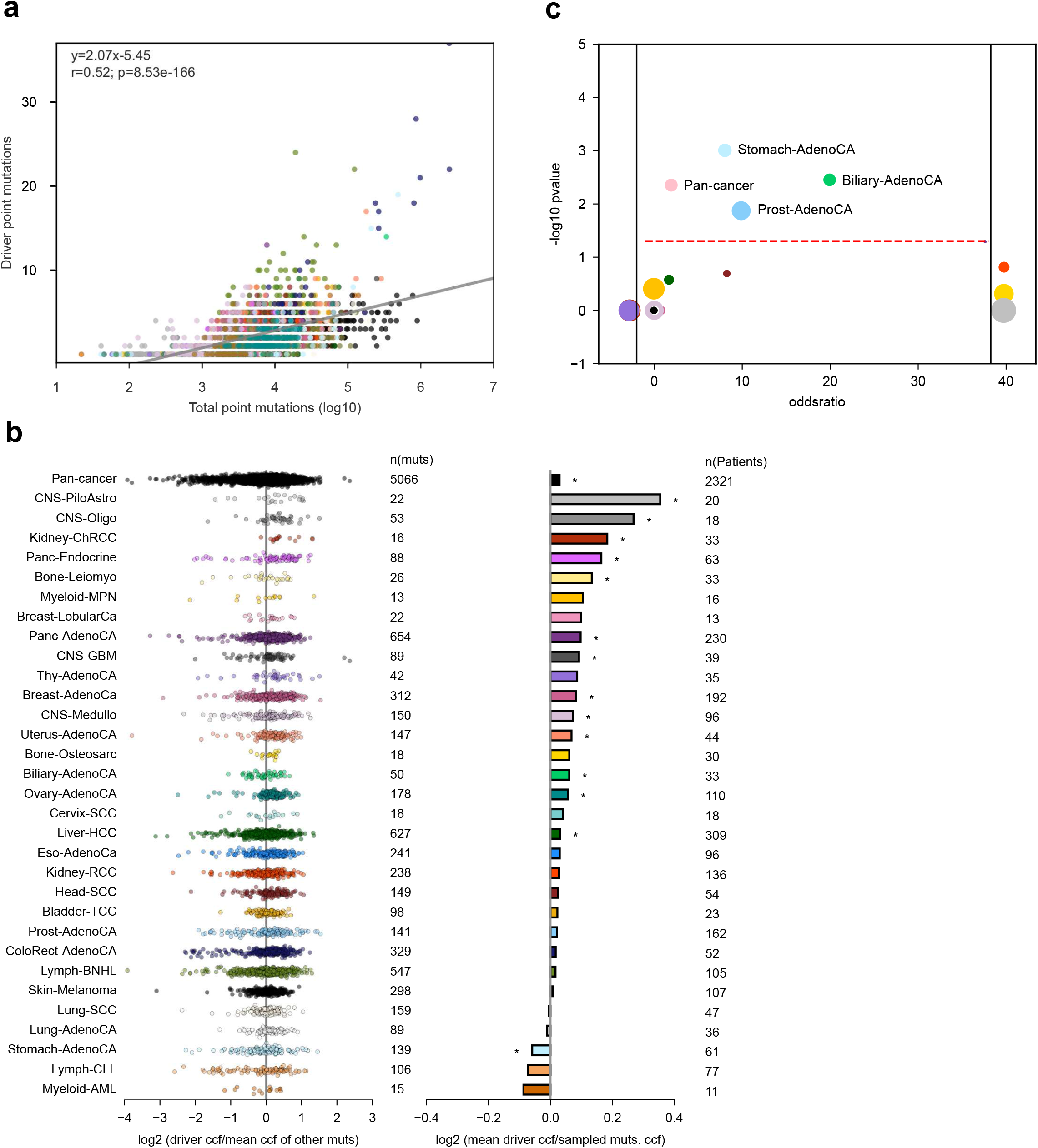
Features of the catalog of driver point mutations. (a) Correlation between the number of driver point mutations in tumors and their mutational burden (in logarithmic scale). There is a significant positive correlation between these two quantities, showing that tumors with more mutation burden carry more driver point mutations. Nevertheless, the number of driver point mutations increases slower that the total number of point mutations in PCAWG tumors: the slope of the regression between the number of driver point mutations and the logarithm of the total number of point mutations is approximately 2. (b) Driver point mutations overall have significantly higher Clonal Cell Fraction (CCF) than other mutations observed in tumors. The left panel shows the comparison of the CCF of each driver mutation to the average CCF of other mutations in the same tumor, and the right panel shows the comparison of the average CCF of driver mutations to the average CCF of driver mutations in each cohort. The cohorts with significantly (P<0.01) higher or lower CCF of driver mutations than randomly sampled mutations are highlighted (*). (c) In some cohorts, patients with no identified driver point mutation are significantly enriched (Fisher’s p-value<0.05) for tumors with insufficient coverage (estimated via the number of reads per clonal copy, or NRPCC) to call point mutations reliably (NRPCC<5). In the graph, each cohort is represented as a circle, located according to the odds-ratio and the p-value of the Fisher’s test. The size of eah circle is proportional to the fraction of patients with no driver point mutations in the cohort. Cohorts with no drivers in patients with sub-threshold NRPCC (infinite odds-ratio) appear at the right margin of the graph; cohorts with no sub-threshold NRPCC patients (undetermined odds-ratio) appear at the left margin of the graph.

**Extended Data Figure 4.**
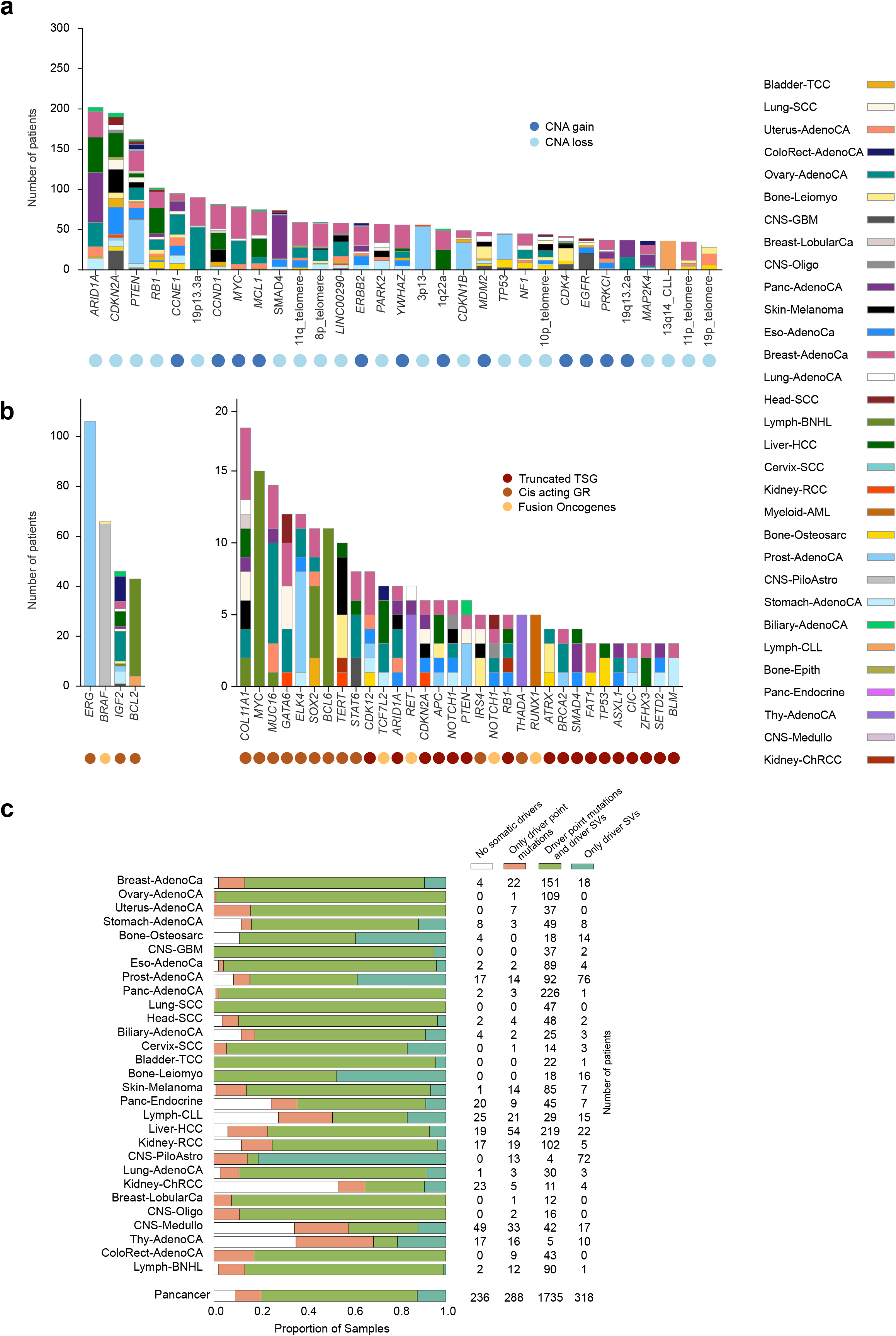
Driver SVs in the panorama. (a) Most recurrent driver SCNA gains and losses in the PCAWG cohort. The frequency of each event across cohorts is represented as a row-stacked barplot using the same color legend for tumor types as in previous figures. The names of the events correspond to the driver gene within the event or to the chromosomal band the event overlaps with if it contains no known driver gene. The circles below the bar plot denote the nature of each driver as SCNA gain or loss. (b) Coding genes involved in the most recurrent driver genomic rearrangements in the PCAWG cohort. Note that events involving several partners of translocation may have been added to produce the total count of events for each gene. The frequency of the events involving each gene across the cohort of each cancer type is represented as a row-stacked bar plot using the same color legend for tumor types as in previous figures. The circles below the bar plot characterize each event as truncated tumor suppressor gene, fusion oncogene or cis-activating rearrangement. (c) Distribution of patients with somatic driver mutations, driver SVs or both types of driver mutations across cohorts and across the whole PCAWG. This graph complements the view in Fig. 3c of the paper on the relative contribution of somatic point mutations and SVs to tumorigenesis. Cohorts are sorted following the same order as the violinplots in Fig. 3c.

**Extended Data Figure 5.**
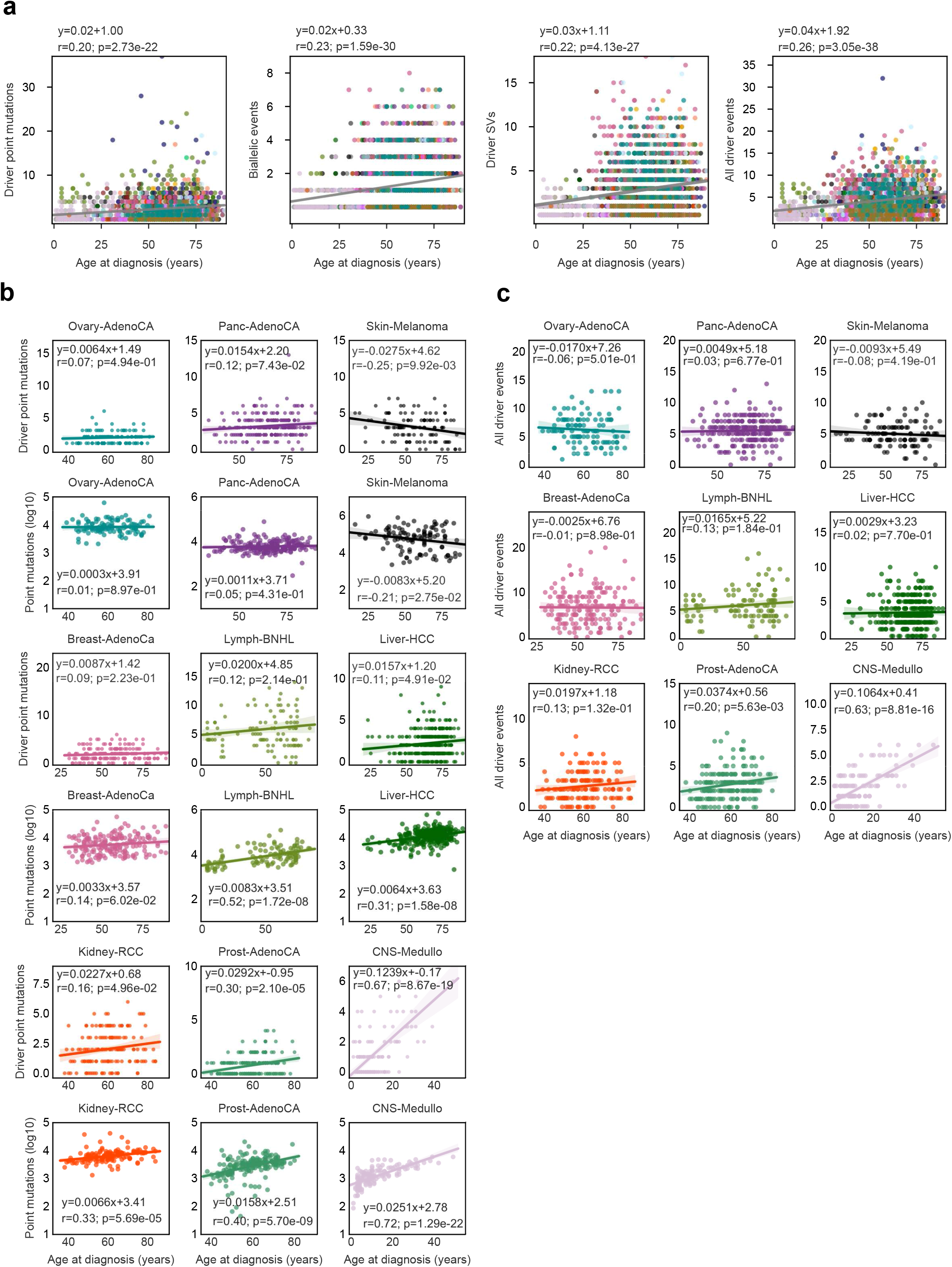
Correlation between the number of driver mutations in tumors and the patients’ age. (a) Correlation between the number of driver point mutations in tumors and patients’ age across the PCAWG cohort. Tumors of different cancer types are represented following the same color convention as throughout the manuscript. From left to right the panels represent the correlation between i) the number of driver point mutations; ii) the number of driver SVs; iii) the number of biallelic events; and iv) the total number of driver events and the age of diagnosis of the tumor. The number of all types of mutations correlates with the age of patients at diagnosis, suggesting that driver mutations accumulate with the progression of tumors, and therefore supporting the idea that our calculation of mean number of driver mutations across the cohort may be higher than the actual minimum number of driver events required to turn a cell malignant. (b) Nevertheless, the correlation between the number of different types of mutations in tumors and patients’ age across selected cancer cohorts (those with at least 100 patients) show striking differences between malignancies. (Odd-number rows present graphs with the correlation between the number of driver point mutations in tumors and the age of diagnosis across three cancer types, and pair-number rows present equivalent graphs with the correlation between the total number of point mutations (in log scale) in tumors and the age of diagnosis.) For certain tumor types (such as prostate adenocarcinomas and medulloblastomas) the mutation burden grows exponentially and the number of drivers grows linearly with the age of diagnosis of the tumor. For some malignancies in this group, such as medulloblastomas, the differences may be related with different courses of the disease in its pediatric and adult presentations. On the other hand, for some cancer types there is no discernible correlation between the total number of point mutations or the number of drivers and the patient’s age (see, for example melanomas and pancreatic adenocarcinomas). This could mean that these tumors tend to be diagnosed mostly in later stages independently of patients’ age. While these differences may reflect actual dissimilarities in the course of these diseases, these results must be taken with caution due to the small sample sizes of these cohorts. (c) Correlations between the total number of driver mutations and the age of diagnosis of the tumor across the same cohorts represented in panel (b).

**Extended Data Figure 6.**
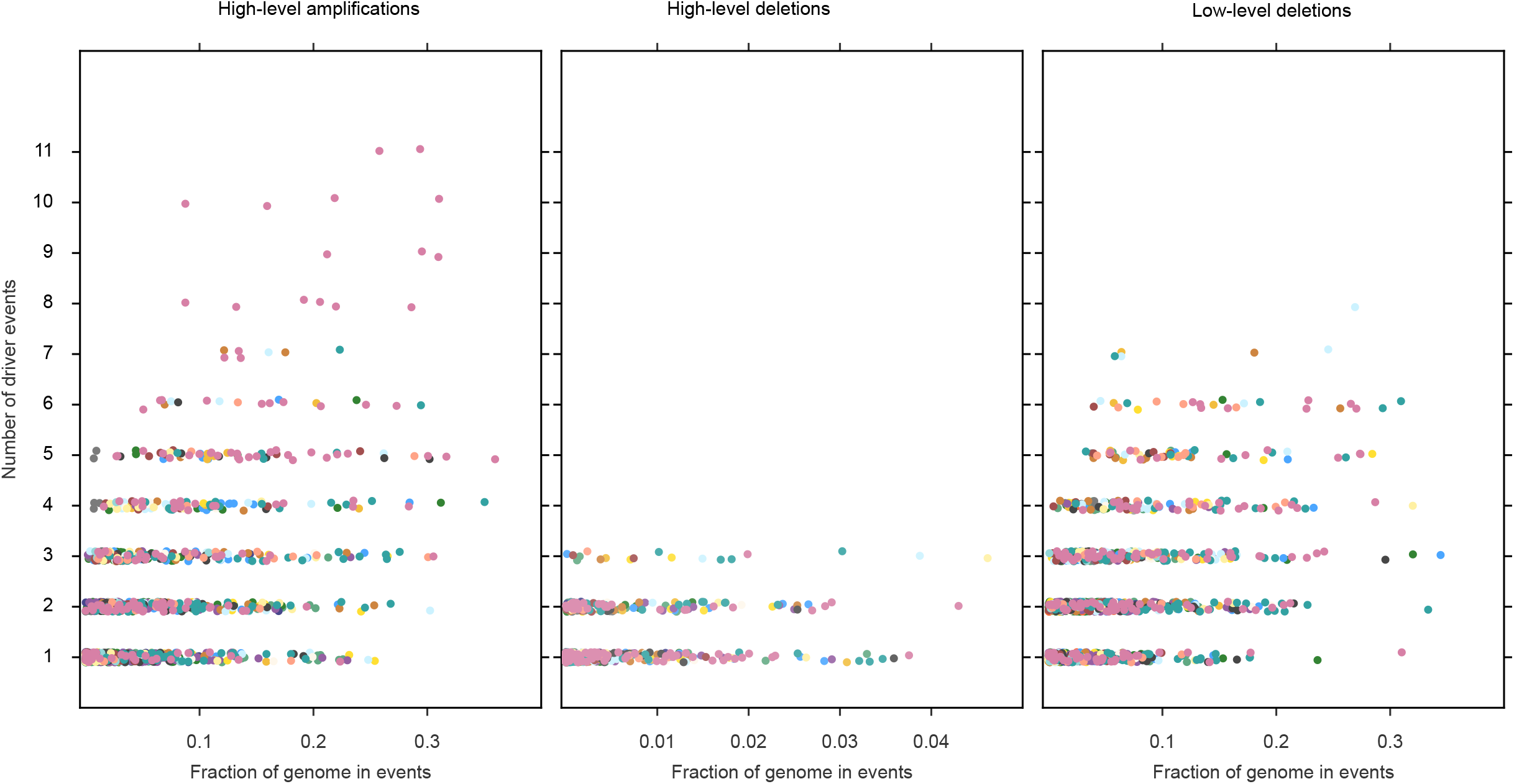
Driver SCNAs across tumors with varying focal SCNAs burden. The graph represents the fraction of the genome of tumors with different number of driver SCNA events that is involved in different types of focal SCNA events. Each dot represents a tumor, colored following the same legend to represent tumor types as in previous figures. The three panels represent the relationship between the number of driver high-level SCNA gains (left), driver high-level SCNA losses (center), and driver low-level SCNA losses (right), with the fraction of the genome involved in focal events of the same nature in each case. The figure demonstrates the lack of a trend towards an increase of the number of driver events in tumors with higher fractions of the genome involved in focal events.

**Extended Data Figure 7.**
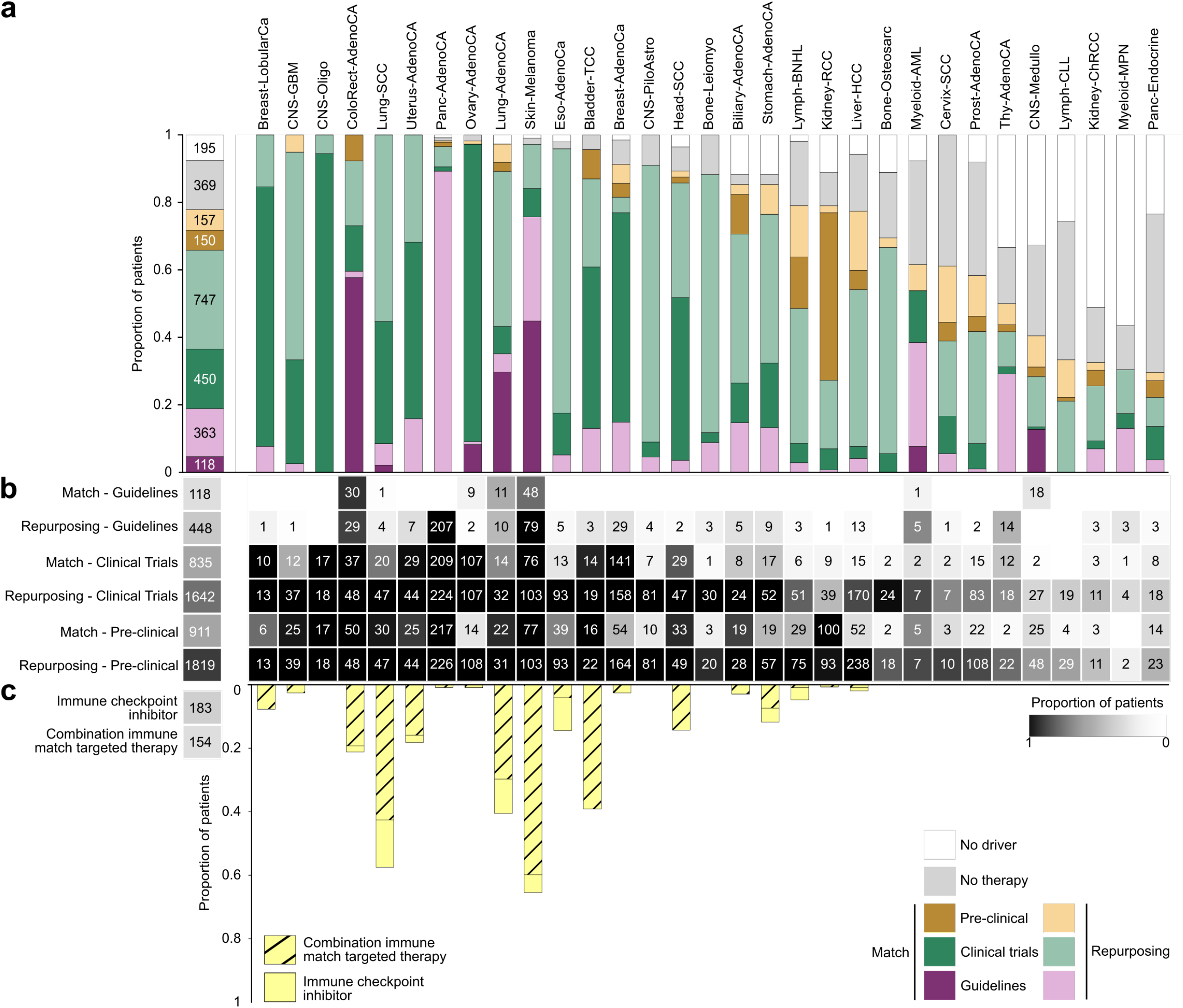
Landscape of anti-cancer actionability of the PCAWG cohort. (a) The landscape of all anti-cancer therapeutic opportunities uncovered by the panorama of driver mutations is represented as a row-stacked bar plot containing the fraction of tumors in each cancer type –and in the pan-cancer cohort– with therapeutic opportunities of different type (color-coded). These include opportunities arising from alterations that are known biomarkers of the response of a tumor to a drug (strong colors) and opportunities of repurposing of these drugs to different tumor types or different types of alterations (faded colors). In this bar plot, each tumor is only counted once with the targeting opportunity that is closest to the clinical use of the drug. (b) The heatmap represents the fraction of patients with therapeutic opportunities of each category (rows) in each tumor type. Here, all opportunities to target each tumor are accounted for. (c) Furthermore, we identified the tumors with a mutation burden above a threshold reported to accurately predict the effectiveness of immune checkpoint therapies to treat a variety of tumors. We found that 7% of tumors in the cohort including 65% of melanomas and 57% of lung squamous cell carcinomas tumors might in principle benefit from these therapies. The fraction of tumors of each cohort susceptible of responding to either immune checkpoint inhibitors or a combination of these drugs with classic anti-cancer drugs considered in the landscape described above is represented as a rowstacked bar plot.

**Extended Data Figure 8.**
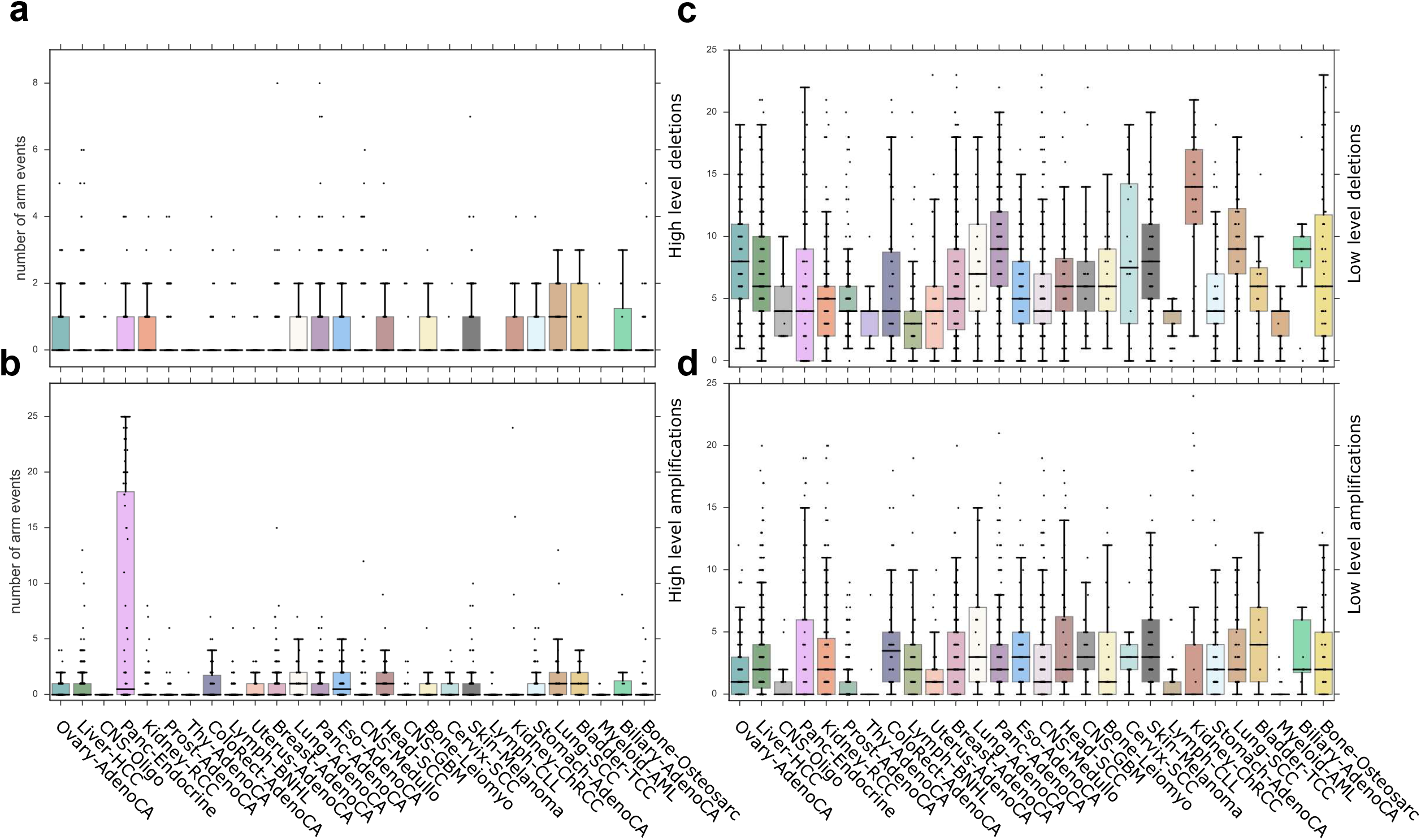
The landscape of arm-level SCNAs across tumors. The graphs show the distribution of the number of arm-level SCNA events observed in each cohort. (a) Distribution of the number of arm high-level SCNA loss events in each cohort. (b) Distribution of the number of arm high-level SCNA gain events in each cohort. (c) Distribution of the fraction of the number of arm low-level SCNA loss events in each cohort. (d) Distribution of the number of arm low-level SCNA gain events in each cohort. The boxplots representing the distribution of events in tumors of each cohort follow the same color legend as previous figures.

**Extended Data Figure 9.**
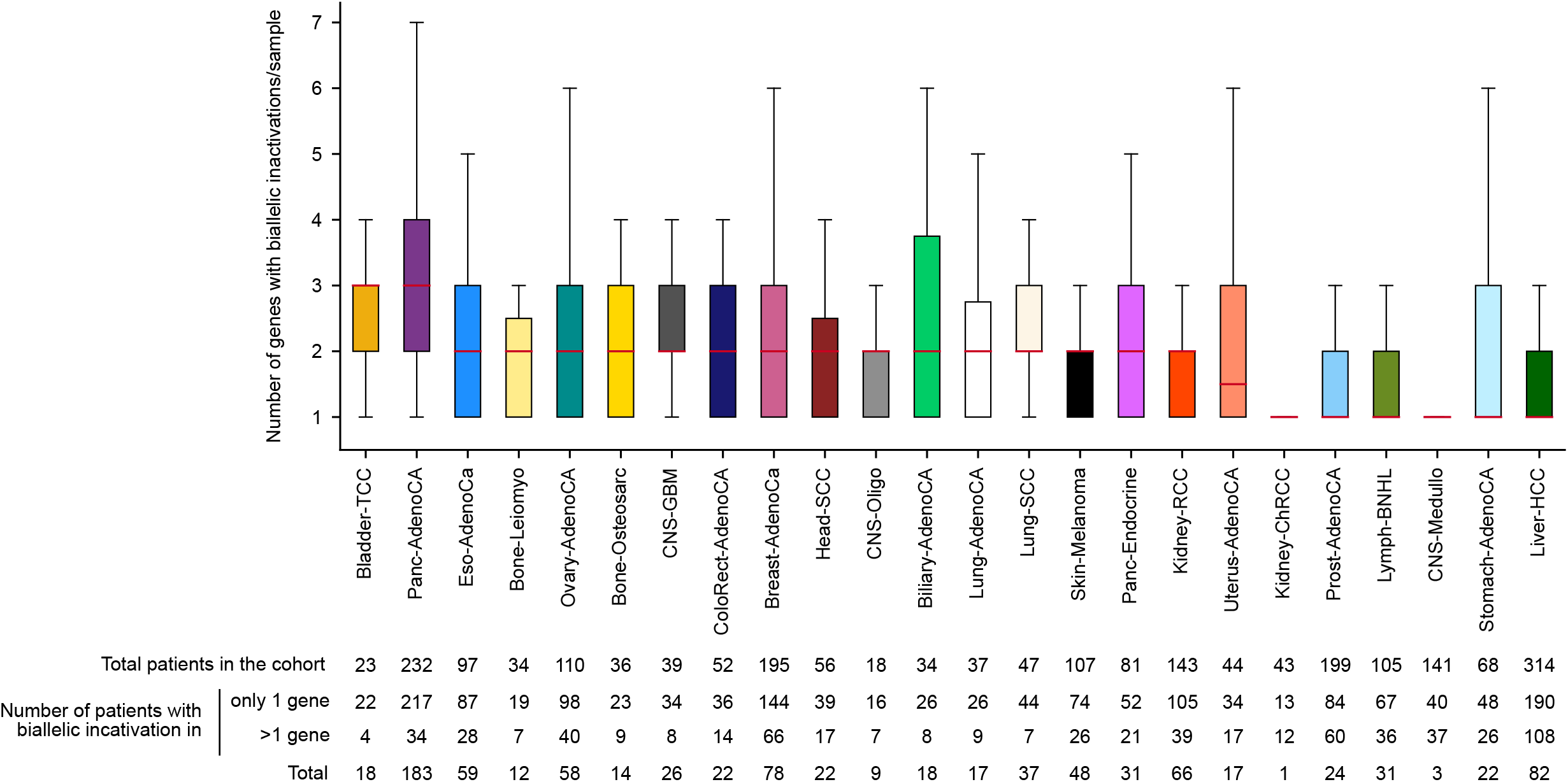
Distribution of biallelic inactivation of tumor suppressor genes. (Number of genes with biallelic inactivation per patients in each cohort. Only cohorts with at least 10 patients affected by one or more biallelic inactivation are included.

**Extended Data Figure 10.**
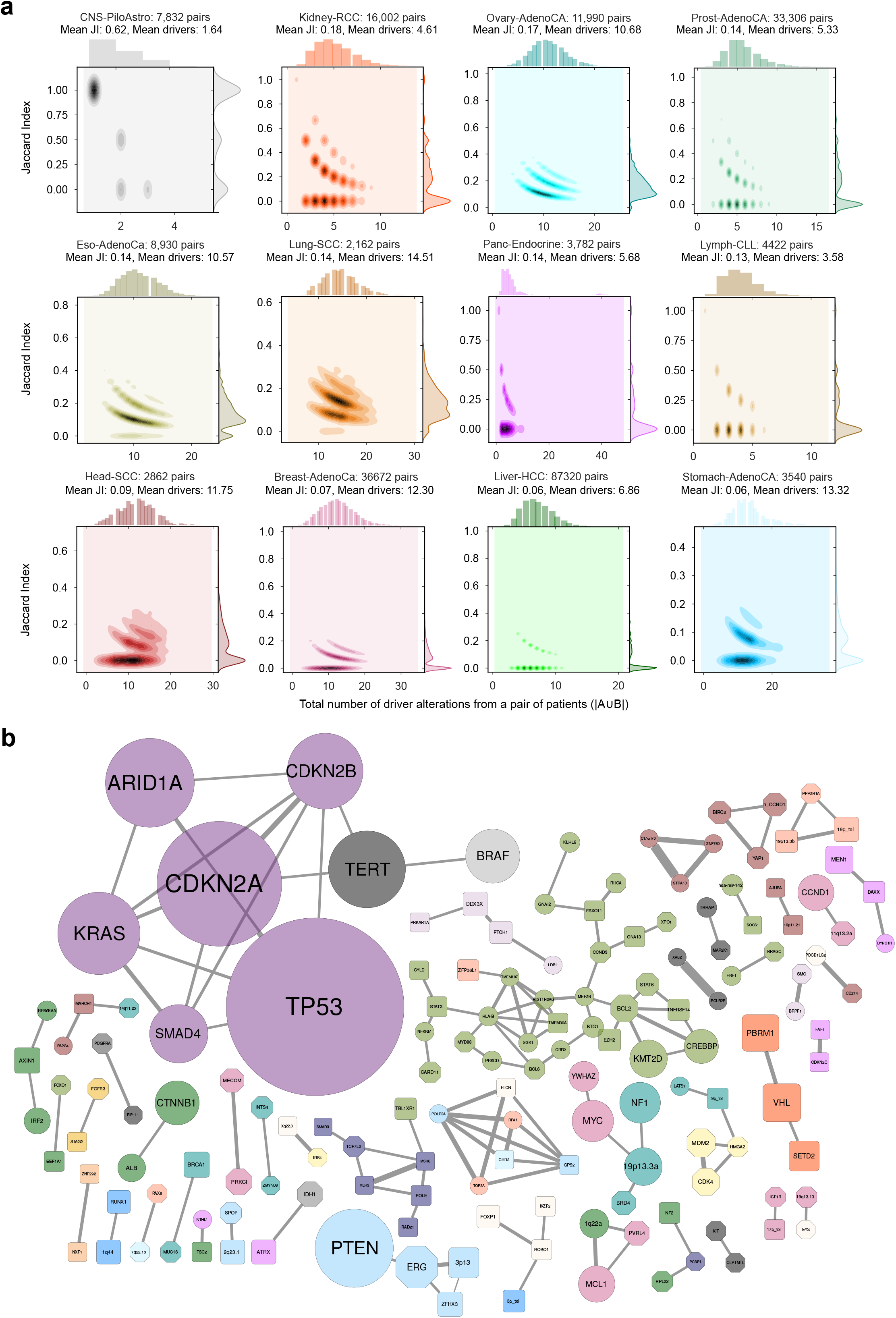
Combinations of drivers across PCAWG tumors. (a) The joinplots with the (smoothed) distributions of Jaccard index (right side) and number of patients in the union (top) and the 2D distribution of both variables across all pairs of patients of 12 tumor types represented in the heatmap of Figure 5a. (b) Network representation of all pairs of GEs with a Jaccard index of overlap of patients with driver mutations above 0. 1. Edges in this network imply high co-occurrence (sharing mutated patients above the Jaccard=0.1 threshold), with their width proportional to this Jaccard index. The size of the nodes (and the font of the gene names) is proportional to the frequency of mutation of the gene, and their color corresponds to the cohort with the highest fraction of driver mutations. The shape of the nodes represent the gene’s mode of action in tumorigenesis.

**Supplementary Table 1. The Compendium of Mutational Driver Elements**

**Supplementary Table 2. Annotated driver point mutations**

**Supplementary Table 3. The panorama of driver mutations in PCAWG**

